# CDK9 interacts with a RanGTP-NEMP1-Importin-β complex to regulate erythroid enucleation

**DOI:** 10.1101/2025.02.03.636174

**Authors:** Lucas M. Newton, Krystle Y. B. Lim, Donia Y. Abeid, Christina B. Wölwer, Chad J. Johnson, Sarah M. Russell, Edwin D. Hawkins, Patrick O. Humbert

## Abstract

Erythroid enucleation is the final stage of erythroid terminal differentiation and involves the separation of an orthochromatic erythroblast into two daughter cells; a pyrenocyte containing the extruded nucleus, and a reticulocyte that will become a red blood cell. Our previous work identified CDK9 as a regulator of erythroid enucleation that appears to act independently of its known role in regulating RNA polymerase II transcription, suggesting the potential for a new CDK9 role. Using a co-immunoprecipitation and mass spectrometry approach, we identified the interactome of CDK9 in differentiating erythroblasts. We show that CDK9 interacts with a RanGTP-NEMP1-Importin-β complex during erythroid terminal differentiation, and inhibition of importin-β in erythroblasts blocks erythroid enucleation. Using imaging analysis and functional assays of enucleating erythroblasts, we show that CDK9 and importin-β co-locate at a critical site of activity opposite to the nucleus before nuclear extrusion and we describe a novel finding that physically links CDK9 and importin-β activity prior to CaM/Ca^2+^ signalling and subsequent F-actin activity to achieve enucleation.

**Key Points:**

- Importin-β physically interacts with CDK9 in erythroid cells and is a novel regulator of erythroid enucleation
- CDK9 and importin-β are required upstream of CaM/Ca^2+^ signalling to enable nuclear extrusion

## Introduction

Erythroid enucleation is the asymmetrical separation of the future erythrocyte from its nucleus and is the final stage of erythropoiesis before the developing erythroblast enters the peripheral blood stream (Palis, 2014). Enucleation is a rate limiting step in efforts to produce red blood cells *ex vivo* for transfusion medicine (Christaki et al., 2019; Gallego-Murillo et al., 2022), necessitating a clearer understanding of its regulation. Several recent reviews describe what is currently known of the molecular and cellular steps required for enucleation (Menon & Ghaffari, 2021; Moras et al., 2017; Newton et al., 2024). At a cellular level, chromatin compaction begins at the polychromatic erythroblast stage, one division prior to the orthochromatic erythroblast, which subsequently undergoes nuclear polarisation and nuclear extrusion; the anucleate cell, termed a reticulocyte, continues developing in the bone marrow for a short time while it undergoes further morphological changes and organelle clearance before moving into the circulation (Menon & Ghaffari, 2021; Moras et al., 2017; Newton et al., 2024). Importantly, the extruded nucleus, termed pyrenocyte, is contained within a separate cell membrane and enclosed in a thin layer of cytoplasm, suggesting the mechanism is not purely exocytosis of nuclear material (Yoshida et al., 2005). Enucleation has been ^comp2ared^ to both cytokinesis, and the apoptosis-driven denucleation events observed in lens epithelia and keratinocytes (Rogerson et al., 2018; Wride, 2011), however the mechanisms that underpin enucleation appear quite distinct from either process (Newton et al., 2024). The molecular regulation of this unique cellular process is poorly understood, but we recently made the surprising finding that CDK9 is essential for erythroid enucleation (Wölwer et al., 2015). Interestingly, our findings also suggested that CDK9 did not act through the canonical RNA Pol II-dependent pathway (Anshabo et al., 2021; Paparidis et al., 2017; Wölwer et al., 2015). Despite its importance, the precise mechanisms that govern enucleation remain relatively poorly understood.

Cytoskeletal elements such as F-actin and non-muscle myosin IIB (NMIIB) are required for the nuclear extrusion process, and while the formation of a contractile actomyosin ring (CAR) has been proposed (Ji et al., 2008; Konstantinidis et al., 2012; Koury et al., 1989; Li et al., 2017), other models suggest that enucleation is achieved primarily through the activity of an actin-mediated enucleosome and by membrane reorganisation by trafficking of endocytic vesicles (An et al., 2021; Keerthivasan et al., 2010; Liang et al., 2015; Nowak et al., 2017). Calcium signalling through the calmodulin (CaM) pathway is critical to achieve enucleation following uptake of extracellular calcium prior to nuclear extrusion (Wölwer et al., 2016). This is likely due to the ability for CaMKII to modulate the actin cytoskeleton by altering F-actin cross-linking and the rate of F-actin polymerisation (Hoffman et al., 2013; Ohta et al., 1986; Sanabria et al., 2009).

Although many of the above factors are well established, the use of standard genetic approaches to identify new regulators of enucleation can be problematic in terms of their interpretation. This is because of the possible indirect effects of genetically mediated perturbation, when using CRISPR or RNAi approaches on upstream processes in terminal differentiation, which can indirectly result in reduced downstream enucleation. To circumvent this, we have developed and used a chemical genetics approach to identify novel regulators of enucleation. We initially identified Cyclin-dependent kinase-9 (CDK9) as a novel regulator of erythroid enucleation (Wölwer et al., 2015). CDK9 is predominantly known for its role as the catalytic subunit of the positive-transcription and elongation factor-B (P-TEFb), which alongside its cyclin partner cyclin T1 (CycT1), cyclin T2a/b (CycT2a/b) or cyclin K, plays a fundamental role in the regulation of RNA polymerase II (RNA Pol II) transcriptional activity (Paparidis et al., 2017; Qin et al., 2017). However, our previous study had shown that blocking the activity of RNA Pol II in late erythropoiesis does not arrest enucleation (Wölwer et al., 2015), suggesting a new role for CDK9 in enucleation independent of RNA Pol II.

Here, using mouse and human erythroid culture models, we characterised the role of CDK9 in erythroid enucleation. We conducted imaging analysis of enucleating erythroblasts and found that CDK9 and Cyclin T1 become more cytosolic and localise nearby F-actin prior to nuclear extrusion. Additionally, we show that CDK9 localisation is dependent on F-actin activity. To identify the pathways CDK9 may be regulating in enucleation, we carried out a proteomics screen to identify relevant CDK9 interactors during erythroid enucleation. We show that CDK9 associates with a RanGTP-NEMP1-Importin-β complex and that CDK9 and importin-β function upstream of calcium signalling and F-actin activity prior to nuclear extrusion. Our study provides new insights into the function of CDK9 in enucleation, and identifies a key link between the new regulator of enucleation Importin-β and previously described regulators CDK9 (Wölwer et al., 2015) and NEMP1 (Hodzic et al., 2022). As CDK9 inhibitors are being tested for clinical use against a variety of cancers, our study also highlights potential side-effects of CDK9 inhibitors on erythropoiesis and enucleation.

## Material and Methods

### Materials

Antibodies, inhibitors and other reagents are listed in supplemental Table 1. pBABE-Flag-Cdk9-IRES-eGFP, pBABE-Flag-Cdk9-T186A-IRES-eGFP and pBABE-Flag-Cdk9-D167N-IRES-eGFP were gifts from Andrew Rice (Addgene plasmid #28096, RRID: Addgene_28096; Addgene plasmid #28097, RRID: Addgene_28097; Addgene plasmid #28098, RRID: Addgene_28098). pBABE GFP was a gift from William Hahn (Addgene plasmid #10668, RRID: Addgene_10668).

### Isolation of orthochromatic erythroblasts

All animal procedures were approved by the La Trobe University Animal Ethics Committee. Splenic orthochromatic erythroblasts were isolated from C57Bl/6 mice between 6-14 weeks of age by phenhylhydrazine treatment and flow cytometry as previously described (Wölwer et al., 2015). Briefly, we sorted for viable (propidium iodide negative) Ter119^+^CD44^low^ cells, which have been previously validated to be primarily orthochromatic erythroblasts (Wölwer et al., 2015). In normal conditions, we regularly observed ∼60% enucleation 5 hours after flow cytometric sorting, which remained stable for up to 16 hours.

### HUDEP-2 and BEL-A cell culture

HUDEP-2 and BEL-A cells were cultured as previously described (Trakarnsanga et al., 2017).

### Flow cytometry

Enucleation of mouse orthochromatic erythroblasts was quantified using a FACSymphony A3 (Becton Dickson; Franklin Lakes, NJ) running FACS Diva software (Becton Dickson; Franklin Lakes, NJ) and analysed using FlowJo v10.8.1 (Becton Dickson; Franklin Lakes, NJ). Percentage of enucleation was obtained by dividing the number of enucleated cells (Ter119^+^ Hoechst ^-^) by the sum of enucleated cells and erythroblasts (Ter119^+^ Hoechst^+^) and multiplying by 100. Propidium iodide was used to eliminate dead cells from the analysis.

### Cytospins

6 × 10^4^ cells were centrifuged onto slides at 320 rpm for 4 minutes using a Cytospin 4 Centrifuge (Thermo Fisher; Waltham, MA). Slides were air dried prior to fixation in methanol and subsequently stained using RapidDiff as per the manufacturer’s guidelines. HUDEP-2 and BEL-A enucleation rates, and cell morphologies were quantified manually by images captured using an Olympus IX81 microscope using 100x/1.4NA oil objective running cellSens Dimension software (Evident/Olympus Life Sciences; Tokyo, Japan).

### Immunofluorescence confocal microscopy

Cells were prepared for immunofluorescence microscopy as previously described (Smith et al., 2018) with minor alterations. Briefly, cells were collected by centrifugation following overnight incubation +/- inhibitors, washed in PBS and immediately fixed in 4% paraformaldehyde in PBS overnight at 4°C. Cells were subsequently washed 3 times in PBS and permeabilised in PBS containing 0.3% Triton X-100 for 15 minutes at RT, before blocking with PBS containing 3% BSA, 1% donkey serum for 2 hours at RT. Cells were incubated with primary antibodies in blocking buffer overnight at 4°C, washed 3 times in PBS and incubated with secondary antibodies and 1 µg/mL DAPI in blocking buffer for 2 hours. Cells were washed 3 times in PBS before centrifugation onto poly-D-lysine treated microscopy slides using a cytospin centrifuge, and subsequently mounted in ProLong Gold antifade reagent. Slides were imaged using a Zeiss LSM800 confocal microscope using 63x/1.4NA oil objective running Zeiss Zen Blue software (Zeiss; Oberkochen, Germany). Nuclear to cytoplasmic signal ratios were calculated using Fiji (ImageJ) (Schindelin et al., 2012). Our custom nuclear to cytoplasmic ratio ImageJ macro is available on GitHub.

### Retroviral transduction of HUDEP-2 cells

Retrovirus was generated by calcium phosphate transfection of HEK293T cells as previously described (Jordan et al., 1996). At 12 hours post-transfection, transfection media was replaced with HUDEP-2 expansion media. Viral media was collected after 24 hours, and HUDEP-2 cells were seeded at 3 x10^5^ cells/mL in viral growth media for 24 hours. Transduced cells were isolated using FACS Aria Fusion (Becton Dickson; Franklin Lakes, NJ) running FACS Diva software (Becton Dickson; Franklin Lakes, NJ) by sorting for GFP^+^ cells.

### CDK9 co-immunoprecipitation, mass spectrometry and protein database search

Detailed methodology is available in the Supplementary Methods. Co-immunoprecipitation studies were performed using the Pierce Classic Magnetic IP/Co-IP Kit (Thermo Fisher; Waltham, MA) as per the manufacturer’s instructions with minor modifications. Protein extracts were quantified using the DC Protein Assay (BioRad; Hercules, CA). Protein samples were dried and desalted before liquid chromatography tandem mass spectrometry (LC MS/MS) using a Thermo Ultimate 3000 RSLC nano UHPLC system and analysed on a Thermo Q-Exactive HF Orbitrap mass-spectrometer (Thermo Fisher Scientific, Waltham, MA). On-bead trypsin digestion was carried out as previously described (Antonicka et al., 2020).

The protein database searches were conducted using Sequest HT search engine via the Proteome Discoverer 2.4 software suite (Thermo Fisher Scientific, Waltham, MA). Spectra were matched against the Homo sapiens reference proteome downloaded from Uniprot (The UniProt Consortium, 2023). Proteins that were present in all three replicates with a minimum 3-fold enrichment compared to the IgG controls were considered relevant. Results, including generation of Venn diagrams and heatmaps, were analysed using FunRich v3.1.4 (Fonseka et al., 2021). Interaction and gene ontology (GO) analysis was performed using the STRING database (Szklarczyk et al., 2023). Original protein database search results will be made available on HUPO depository.

### Western blot

Cell lysates were isolated using NETN lysis buffer (250 mM NaCl, 5 mM EDTA (pH 8.0), 50 mM Tris-HCl (pH 8.0), 0.5% NP-40 in distilled water) containing Complete mini protease inhibitor (Roche; Basel, Switzerland) and PhosSTOP phosphatase inhibitor (Roche; Basel, Switzerland). Protein levels were quantified using a DC protein assay (BioRad; Hercules, CA), denatured using SDS loading buffer and subsequently resolved on standard sodium dodecyl-sulfate polyacrylamide gels before western transfer onto polyvinylidene difluoride membrane (Merck Millipore; Burlington, MA). Membranes were blocked in 3% BSA in 0.2% Tween-20 in PBS for 1 hour, then incubated overnight with primary and secondary antibodies respectively. Membranes were visualised using an Odyssey infrared imager (LiCor Biosciences; Lincoln, NE). Antibodies and concentrations used are listed in Supplementary Table 1.

### Statistical analysis

Statistical calculations and analysis were performed using GraphPad Prism version 9.2.0 (GraphPad Software, San Diego, CA). Statistical differences were analysed using ANOVA tests, with Dunnett’s or Tukey’s tests for multiple comparisons. Detailed statistics are described in the figure legends. n = number of cells counted, or number of replicates. P-values below 0.05 were considered statistically significant.

## Results

### Phosphorylated CDK9 (Thr186) and Cyclin T1 become cytoplasmic and localise near F-actin during nuclear extrusion

To better understand how CDK9 might regulate enucleation, we investigated the localisation of CDK9 and key binding partner cyclin T1 during erythroid differentiation and enucleation. We used mouse and human erythroid *in vitro* systems to compare, for two stages of enucleation **(Figure 1A)**, the localisation and phosphorylation (Thr186, to reflect CDK9 activation) of CDK9. In mouse orthochromatic erythroblasts p-CDK9 was observed in smaller clusters within the nucleus during the nuclear condensation phase, before localising to larger clusters within the cytoplasm that are also found in the future reticulocyte during extrusion; these clusters also seem to localise closely with F-actin during nuclear extrusion **(Figure 1B)**. p-CDK9 and CDK9 signal can be seen to co-localise, however total CDK9 protein is more spread throughout the reticulocyte **(Figure 1B)**. Cyclin T1, the main binding partner and regulator of CDK9, was localised to the cytoplasm of mouse orthochromatic cells during nuclear condensation **(Figure 1C)**.

**Figure 1.**
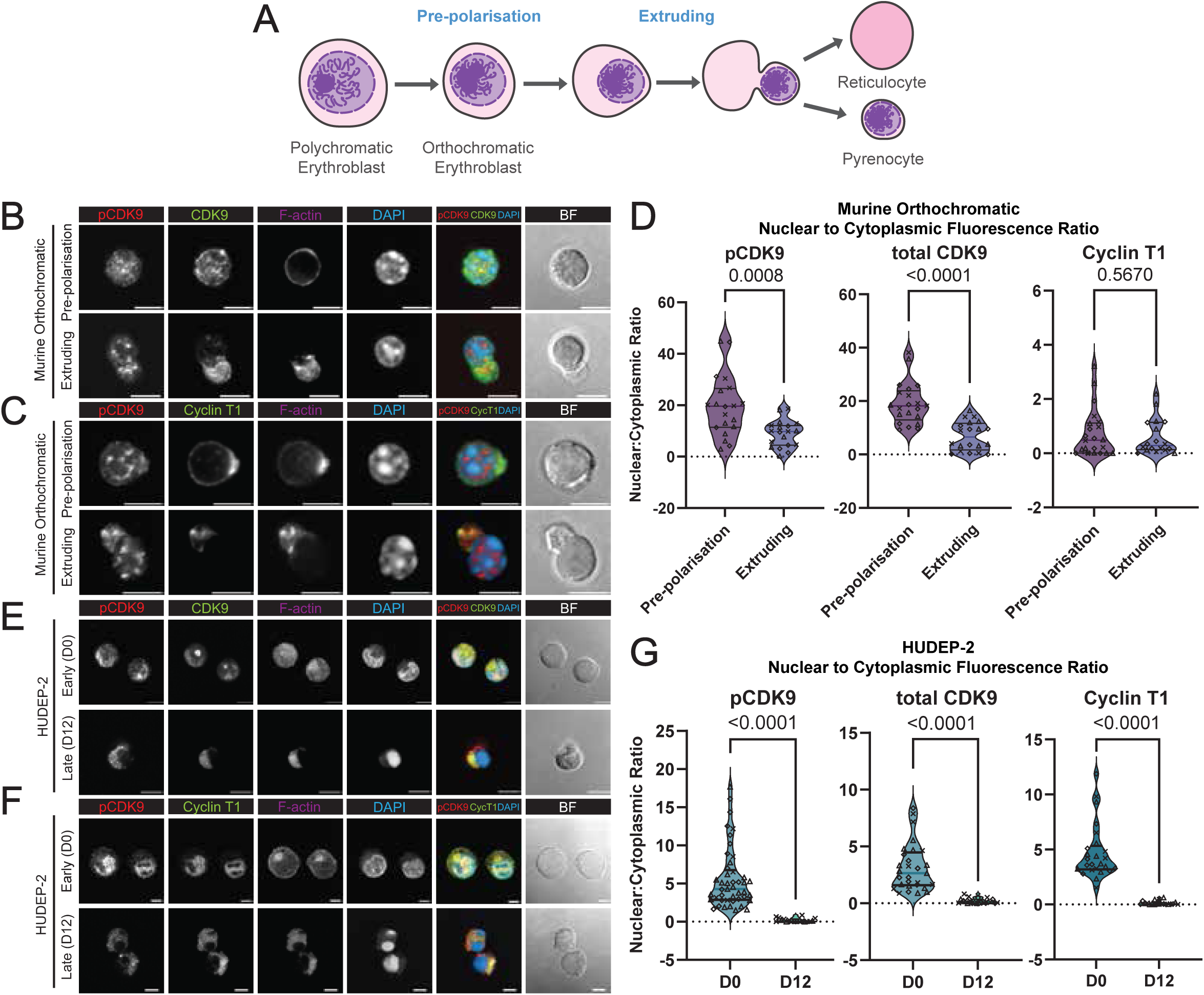
Phosphorylated CDK9 (Thr186) and Cyclin T1 are excluded from the nucleus during nuclear extrusion and localise to the cytoplasm and future reticulocyte following enucleation. **(A)** Diagram depicting the stages of erythroid enucleation described in this study. **(B)** Immunofluorescence confocal microscopy of mouse orthochromatic erythroblasts at the pre-polarisation and extrusion phases of erythroid enucleation stained for phospho-CDK9(Thr186), CDK9 (F-6), phalloidin for F-actin and DAPI for nuclei. Brightfield (BF) images and actin are excluded from the merge. Yellow colour on the merge indicates co-localisation of p-CDK9 and CDK9. All scale bars = 5 μm. **(C)** Immunofluorescence confocal microscopy of mouse orthochromatic erythroblasts at the pre-polarisation and extrusion phases of erythroid enucleation stained for phospho-CDK9(Thr186), Cyclin T1, phalloidin for F-actin and DAPI for nuclei. Brightfield (BF) images and actin are excluded from the merge. Yellow colour on the merge indicates co-localisation of p-CDK9 and Cyclin T1. All scale bars = 5 μm. **(D)** Nuclear to cytoplasmic ratios of p-CDK9 (n = 20 pre-polarisation, 19 extruding orthoblasts), total CDK9 (n = 21 pre-polarisation, 20 extruding orthoblasts) and Cyclin T1 (n = 22 pre-polarisation, 18 extruding orthoblasts). **(E)** Immunofluorescence confocal microscopy of HUDEP-2 cells at early (day 0) and late (day 12) of differentiation stained for phospho-CDK9(Thr186), CDK9 (F-6), phalloidin for F-actin and DAPI for nuclei. Brightfield (BF) images and actin are excluded from the merge. Yellow colour on the merge indicates co-localisation of p-CDK9 and CDK9. All scale bars = 5 μm. **(F)** Immunofluorescence confocal microscopy of HUDEP-2 cells at early (day 0) and late (day 12) of differentiation stained for phospho-CDK9(Thr186), Cyclin T1, phalloidin for F-actin and DAPI for nuclei. Brightfield (BF) images and actin are excluded from the merge. Yellow colour on the merge indicates co-localisation of p-CDK9 and Cyclin T1. All scale bars = 5 μm. **(G)** Nuclear to cytoplasmic ratios of p-CDK9 (n = 45 day 0 (D0), 18 day 12 (D12) HUDEP-2 cells), total CDK9 (n = 29 day 0 (D0), 22 day 12 (D12) HUDEP-2 cells) and Cyclin T1 (n = 25 day 0 (D0), 19 day 12 (D12) HUDEP-2 cells).

Upon measuring nuclear to cytoplasmic fluorescence ratios, we observed a strong shift towards the cytoplasm for p-CDK9 and total CDK9 **(Figure 1D)**. Cyclin T1 remains primarily cytosolic before colocalising with p-CDK9 in clustered regions near F-actin during nuclear extrusion **(Figure 1C & D)**. These findings were recapitulated during enucleation in the human erythroblast cell line HUDEP-2 **(Figure 1E & F)**. Quantifying these images reveals a complete exclusion of p-CDK9, CDK9 and Cyclin T1 in day 12 (D12) differentiated HUDEP-2 cells from the nucleus which appears highly condensed **(Figure 1G)**. Thus, CDK9 remains active in the nucleolus very late into terminal differentiation and subsequently localises with F-actin and cyclin T at the rear of the extruding nucleus before relocating to the future reticulocyte in both mouse and human enucleating erythroblasts.

### Highly selective inhibition of CDK9 activity arrests enucleation at the same stage as F-actin induced enucleation block

Our previous study revealed that pharmacological blockage of CDK9 activity leads to an arrest in enucleation, and together with the observation that CDK9-Cyclin T1 co-localise with F-actin, suggests that CDK9 might regulate the extrusion event. To examine this potential functional significance, we initially compared the effects of CDK9 inhibition with inhibition of actin polymerisation. With the advent of next generation high selectivity CDK9 inhibitors, we first confirmed the effect of specific CDK9 inhibition on enucleating cells in comparison to the F-actin inhibitor cytochalasin D (CytoD), which consistently arrests cells at a late stage of enucleation (Konstantinidis et al., 2012; Wang et al., 2012; Wölwer et al., 2015) **(Figure 2)**. Treatment of orthochromatic mouse erythroblasts **(Figure 2A)**, with CDK9 inhibitors NVP-2 (Olson et al., 2017) **(Figure 2B)** and AZD4573 (Barlaam et al., 2020) **(Figure 2C)** arrested enucleation in a dose dependent manner, with a mean reduction of 51% and 64% against the vehicle (DMSO) control at 10 nM respectively. Phenotype analysis of cytospun cells reveal similar arrest patterns for both inhibitors consistent with the phenotypes reported previously for less specific CDK9 inhibitors (Wölwer et al., 2015), with an increased percentage of cells with polarised or partially extruded nucleus (**Figure 2B, C)** compared to the control **(Figure 2A)**. NVP-2 also reduced enucleation of human cells compared to the vehicle (DMSO) control, with a similar increase in polarised nuclei **(Supp. Figure 1A-1B)**. This data confirms that the action of CDK9 on erythroid enucleation is conserved between mouse and human, and further validates our previous findings by alleviating concerns around inhibitor off-target effects on other CDKs described here (Wölwer et al., 2015).

**Figure 2.**
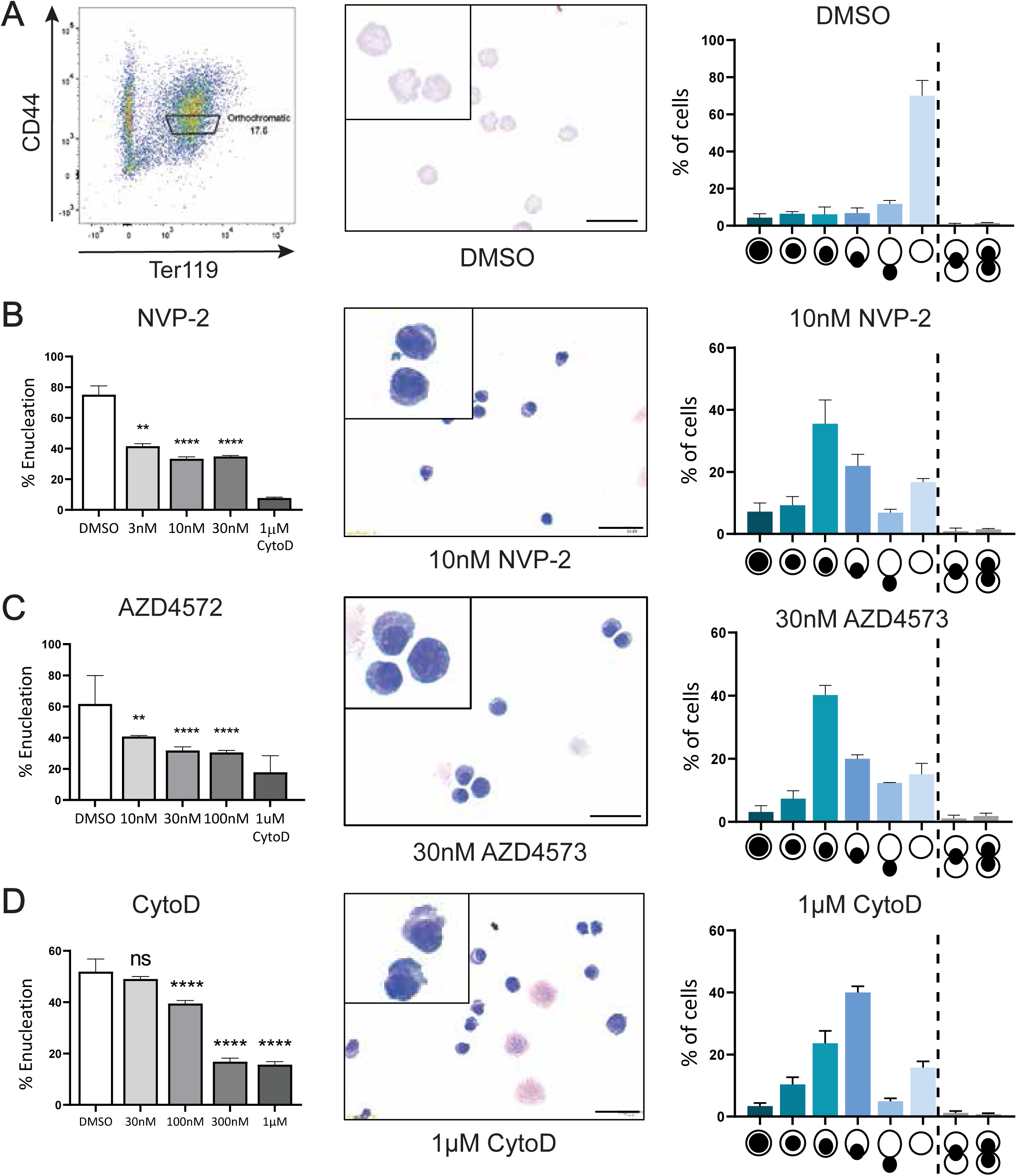
CDK9 inhibitors cause an enucleation arrest that phenocopies cytochalasin D-mediated enucleation blockage. **(A)** Gating strategy used for FACS isolation of orthochromatic erythroblasts from mouse spleen following phenylhydrazine treatment alongside cytospin rapid diff staining and phenotype analysis following 12-hour incubation of isolated cells. **(B)** Quantification of enucleation of mouse orthochromatic erythroblasts following treatment with NVP-2 for 12 hours. DMSO (vehicle control) and cytochalasin D (positive control) are included, in addition to cytospin rapid diff staining and phenotype analysis. n = 4 replicates across 3 independent experiments. (\*\**p* < 0.01, \*\*\*\**p* < 0.0001; one-way ANOVA with Dunnett’s multiple comparisons test). Scale bar = 10µm. **(C)** Quantification of enucleation of mouse orthochromatic erythroblasts following treatment with AZD4573 for 12 hours. DMSO (vehicle control) and cytochalasin D (positive control) are included, in addition to cytospin rapid diff staining and phenotype analysis. n = 4 replicates across 3 independent experiments. (\*\**p* < 0.01, \*\*\*\**p* < 0.0001; one-way ANOVA with Dunnett’s multiple comparisons test). Scale bar = 10 µm. **(D)** Quantification of enucleation of mouse orthochromatic erythroblasts following treatment with cytochalasin D (CytoD) for 12 hours. DMSO (vehicle control) is included, in addition to cytospin rapid diff staining and phenotype analysis. n = 4 replicates across 2 independent experiments. (ns = not significant, \*\*\*\**p* < 0.0001; one-way ANOVA with Dunnett’s multiple comparisons test). Scale bar = 10 µm.

All other methods of chemical inhibition of CDK9 provided similar results. Atuveiclib (BAY 1143572) (Lücking et al., 2017), and the thalidomide-conjugate PROTAC degrader THAL-SNS-032 (Olson et al., 2017) both arrested enucleation compared to the vehicle (DMSO) control **(Supp. Figure 2)**. To compare the effects of CDK9 inhibition to a cytochalasin D (CytoD)-induced late-stage enucleation blockage, we treated mouse orthochromatic erythroblasts to a range of concentrations and quantified the phenotype of cytospins using morphology indicators **(Figure 2)**. Examination of cytospins indicate that CDK9 inhibitors phenocopy a blockage by the actin polymerisation inhibitor cytochalasin D (CytoD) **(Figure 2D)**, with quantitation of morphology indicating that cytochalasin D causes a slightly later arrest compared to CDK9 inhibitors, with more cells showing highly polarised and bulging nuclei **(Figure 2D)**. We conclude that CDK9 must act just prior to the final stages of enucleation, with the observed blockage closely resembling that of F-actin inhibition.

### CDK9 is active upstream of calcium signalling and F-actin polymerisation prior to nuclear extrusion

Our localisation studies showed CDK9 co-localisation with F-actin at late stages of enucleation and suggested that CDK9 may be required just upstream of F-actin and therefore may play a role in regulating the nuclear extrusion event. To confirm the order in which activated CDK9, and F-actin assembly may be required for enucleation, we first examined the reciprocal localisation of activated CDK9 and F-actin in the presence of NVP-2 and cytochalasin D (CytoD) in mouse orthochromatic erythroblasts using immunofluorescence imaging **(Figure 3A)**. Cytochalasin D (CytoD), which consistently arrests the cells with a partially extruded nucleus, resulted in accumulation of p-CDK9 at the site of enucleation **(Figure 3A)**. In addition, p-CDK9 signal is diminished in cells treated with NVP-2, as might be expected **(Figure 3A)**. F-actin appears to accumulate and co-localise with p-CDK9 in cytochalasin D (CytoD) treated cells but appears dispersed near the cell membrane in NVP-2 treated cells, where F-actin would normally localise toward the site of enucleation **(Figure 3A)**. Although these experiments indicate a close relationship between CDK9 and actin in the context of enucleation, they did not clarify the signalling hierarchy between CDK9 and actin.

**Figure 3.**
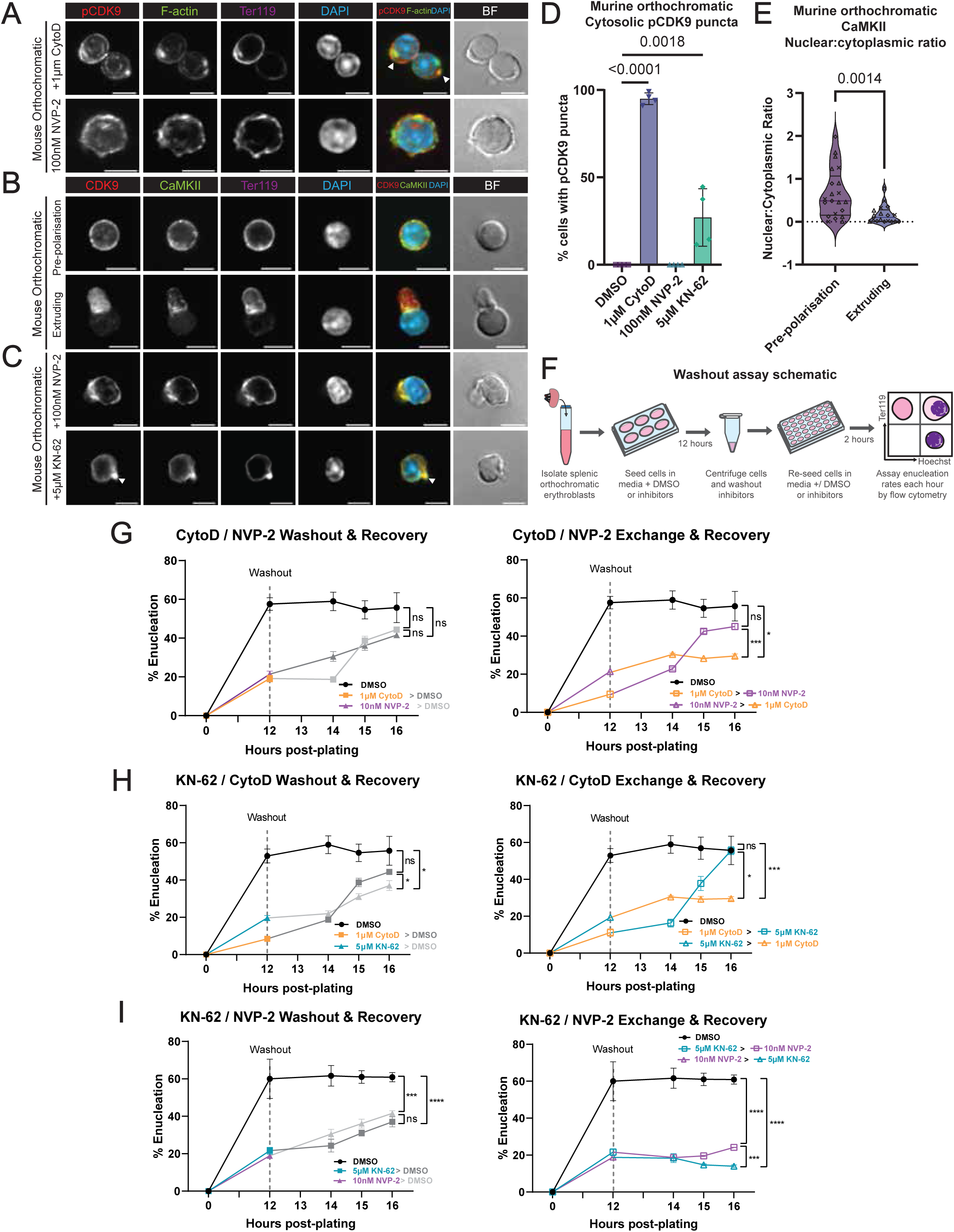
F-actin is required downstream of CDK9 to achieve enucleation, and blocking enucleation with cytochalasin D causes a build-up of active CDK9 that co-localises with F-actin. **(A)** Immunofluorescence confocal microscopy of mouse orthochromatic erythroblasts treated with 1 µM cytochalasin D (CytoD) or 100 nM NVP-2 for 12 hours and stained for phospho-CDK9(Thr186) phalloidin for F-actin, Ter119 and DAPI for nuclei. Brightfield (BF) images and Ter119 are excluded from the merge. Arrows indicate accumulation of CDK9. Yellow colour on the merge indicates co-localisation of p-CDK9 and F-actin. All scale bars = 5 μm. **(B)** Immunofluorescence confocal microscopy of mouse orthochromatic erythroblasts at the pre-polarisation and extrusion phases of erythroid enucleation stained for CDK9 (F-6), CaMKII, Ter119 and DAPI for nuclei. Brightfield (BF) and Ter119 are excluded from the merge. Scale bars = 5 µm. **(C)** Immunofluorescence confocal microscopy of mouse orthochromatic erythroblasts following 12 hours treatment with 100 nM NVP-2 or 5 μM KN-62, stained for CDK9 (F-6), CaMKII, Ter119 and DAPI for nuclei. Brightfield (BF) and Ter119 are excluded from the merge. Arrows indicate accumulation of CDK9. All scale bars = 5 μm. **(D)** Manual scoring of p-CDK9 puncta in orthoblasts following treatment with DMSO (Vehicle; n = 22 cells averaged across 4 experiments), 1µM CytoD (n = 27 cells averaged across 4 experiments), 100nM NVP-2 (n = 21 cells averaged across 4 experiments) and 5µM KN-62 (n = 18 cells averaged across 4 experiments). **(E)** Nuclear to cytoplasmic ratio of CaMKII (n = 21 pre-polarisation, 18 extruding orthoblasts). **(F)** Schematic diagram describing workflow for inhibitor washout and exchange functional assays used in this study. **(G)** Enucleation rates of mouse orthochromatic erythroblasts following inhibitor washout into fresh media containing DMSO (left) or inhibitors (cytochalasin D or NVP-2; right) at 12 hours post-plating. Enucleation rates were measured at 12-, 14-, 15- and 16-hours post-plating. DMSO (black; negative control) is included. Colours change to match either DMSO (grey) or inhibitors (coloured) after washouts. Enucleation rates at hour 16 were statistically compared. n = 5 replicates across 2 independent experiments. (ns = not significant, *\*p* < 0.1, \*\*\**p* < 0.001; two-way ANOVA with Tukey’s multiple comparisons test). **(H)** Enucleation rates of mouse orthochromatic erythroblasts following inhibitor washout into fresh media containing DMSO (first plot) or inhibitors (cytochalasin D or KN-62) at 12 hours post-plating. Enucleation rates were measured at 12-, 14-, 15- and 16-hours post-plating. DMSO (black; negative control) is included. Colours change to match either DMSO (grey) or inhibitors (coloured) after washouts. Enucleation rates at hour 16 were statistically compared (ns = not significant, *p < 0.1, ***p < 0.001; two-way ANOVA with Tukey’s multiple comparisons test). **(I)** Enucleation rates of mouse orthochromatic erythroblasts following inhibitor washout into fresh media containing DMSO (first plot) or inhibitors (KN-62 or NVP-2) at 12 hours post-plating. Enucleation rates were measured at 12-, 14-, 15- and 16-hours post-plating. DMSO (black; negative control) is included. Colours change to match either DMSO (grey) or inhibitors (coloured) after washouts. Enucleation rates at hour 16 were statistically compared (ns = not significant, *p < 0.1, ***p < 0.001; two-way ANOVA with Tukey’s multiple comparisons test).

Calcium signalling is required for effective enucleation, but the positioning of calcium signalling components is not known (Wölwer et al., 2016). We found that in orthochromatic erythroblasts calmodulin kinase II (CaMKII), a key component in CaM/Ca^2+^ signalling, localises primarily in the cytoplasm during nuclear polarisation, and co-localises with CDK9 at the rear of the nucleus during nuclear extrusion **(Figure 3B)**. To better observe the relationship between CDK9 and CaMKII, we examined their localisation in orthochromatic erythroblasts treated with either 100 nM NVP-2 or 5 µM KN-62, which inhibits CaMKII activity (Okazaki et al., 1994) **(Figure 3C)**. CDK9 and CaMKII appear to strongly colocalise under NVP-2 and KN-62 induced blockage, possibly suggesting the existence of a checkpoint dependent on the activity of CDK9 and/or CaMKII. However, we again observed an accumulation of CDK9 near the membrane opposite the polarised nucleus under KN-62 blockage **(Figure 3C)**, suggesting CaMKII signalling may be dependent on upstream CDK9 signalling to facilitate downstream F-actin-mediated nuclear extrusion. We quantified these accumulations of p-CDK9, which we term cytosolic p-CDK9 puncta, which appear in almost all cytochalasin D-treated cells, in some cells treated with KN-62, but never under NVP-2 or DMSO treatment **(Figure 3D)**. Additionally, assessment of nuclear to cytoplasmic ratio of CaMKII reveals a significant shift to the cytoplasm in extruding orthochromatic erythroblasts **(Figure 3E)**.

To assess the order in which CDK9, actin and CaMKII act during enucleation, we applied reversible inhibitors in sequence **(Figure 3G-I)**. We reasoned that the kinetics of recovery would differ depending on whether downstream events were inhibited before or after upstream events. If before, they should not affect recovery, but if after, recovery would be slower. NVP-2 acts to inhibit CDK9 by reversibly blocking the ATP binding site (Olson et al., 2017), and cytochalasin D reversibly disrupts actin polymerisation by forcing actin dimerization (Friederich et al., 1993; Goddette & Frieden, 1986; Stevenson & Begg, 1994). Additionally, KN-62 can reversibly inhibit CaMKII (Okazaki et al., 1994). As NVP-2, cytochalasin D and KN-62 are reversible inhibitors, we used these inhibitors to study the order of events prior to nuclear extrusion. A schematic describing the workflow of these experiments is provided **(Figure 3F)**. Enucleation rates were assessed by flow cytometry following 12 hours incubation with 1 µM cytochalasin D (CytoD) or 10 nM NVP-2 and compared to the vehicle (DMSO) control directly after washout, and at hour 14, 15 and 16 after plating cells **(Figure 3G)**. Enucleation rates following washout of cytochalasin D or NVP-2 into media containing DMSO carrier are shown to recover close to the rate of the DMSO vehicle control at hour 16, confirming the reversibility of both competitive inhibitors **(Figure 3G, Panel 1)**. Exchanging cytochalasin D (CytoD) for NVP-2 at the washout point resulted in a recovery of enucleation like the control, however, exchanging NVP-2 for cytochalasin D (CytoD) resulted in impaired recovery of enucleation to a statistically significant degree compared to both the control and the reverse inhibitor exchange at hour 16 **(Figure 3G, Panel 2)**. These experiments indicate that cells arrested during F-actin polymerisation inhibition by cytochalasin D are not affected by later CDK9 inhibition. Together these results indicate that CDK9 acts upstream of F-actin for nuclear extrusion.

Next, we examined the relationship between actin and CDK9 with CaMKII. Enucleation rates recovered following washout of cytochalasin D (CytoD) and KN-62, with cytochalasin D (CytoD)-treated cells recovering with non-significant variance to the control, and KN-62 washout partially recovering, with enucleation rates still significantly lower than the control at hour 16 **(Figure 3H, Panel 1)**. Inhibition of F-actin (CytoD) after CaMKII signalling (KN-62) results in a complete recovery of enucleation rates by hour 16 compared to the control, but the reverse sequence did not **(Figure 3H, Panel 2)**. This indicates that CaMKII activity is required upstream of F-actin polymerisation prior to nuclear extrusion. Similarly, enucleation rates were assessed following 12 hours incubation with KN-62 or NVP-2 and compared to the vehicle (DMSO) control, where enucleation rates recovered equally in both the KN-62 and NVP-2 treatment to non-significantly different levels compared to the control at hour 16 **(Figure 3I, Panel 1)**. Although washout of KN-62 into NVP-2 allowed for a slight recovery of enucleation, it appeared that no sequence could be conferred, which may suggest that CDK9 and CaM signalling occurs concurrently or independently **(Figure 3I, Panel 2)**. Taken together, these results confirm that both CDK9 activity and CaM signalling is required before F-actin during enucleation.

### CDK9 is associated with a Ran-NEMP1-Importin-β complex in terminally differentiating erythroid cells

Having established that CDK9 acts closely with calcium signalling and upstream of F-actin in its regulation of enucleation, and that our previous work showed that the typical downstream effector of CDK9 being RNA Pol II is dispensable for enucleation (Wölwer et al., 2015), we assessed how CDK9 might modulate actin activity and enucleation in general. We undertook a comprehensive proteomics screen to identify new interactors of CDK9 in the context of erythroid differentiation using CDK9 co-immunoprecipitation mass spectrometry. We used the human HUDEP-2 cell line, as this provided both the sufficient material and the ability to manipulate the system genetically. First, we investigated the interactome of endogenous CDK9 in undifferentiated (Day 0) and differentiated (Day 6) HUDEP-2 cells. We used day 6 differentiated HUDEP-2 cells due to better cell viability and overall protein availability compared to more highly differentiated cells. The original protein database search is available in Supplementary File 2, however, for our analysis we took a stringent cut-off approach, including proteins that were identified in all 3 replicates and ≥3 fold enriched compared to the IgG co-immunoprecipitation controls. 65 unique proteins were identified in undifferentiated HUDEP-2 cells, and 114 unique proteins identified in differentiated HUDEP-2 cells, with 13 proteins identified in both data sets **(Figure 4A)**. We performed a STRING interactome analysis on these 13 proteins. Of note, we identified CDK9 and cyclin T1 (CCNT1) as common to both data sets, validating the presence of a CDK9-Cyclin T1 complex in differentiating human erythroid cells **(Figure 4B)**. Other known interactors of CDK9, including vimentin (VIM) and the DNA-activated protein kinase subunit PRKDC were also found in both data sets (Yang et al., 2015; Zhang et al., 2014). Surprisingly, we also found 5 members of the Ran GTPase nuclear import complex, including nuclear envelope membrane protein 1 (NEMP1/TMEM194A) and importin-β (KPNB1) **(Figure 4B)**. Of note, NEMP1/TMEM194A has recently been described as an essential regulator of enucleation in the mouse (Hodzic et al., 2022), confirming our proteomics screen using human cells can identify functionally relevant proteins for enucleation.

**Figure 4.**
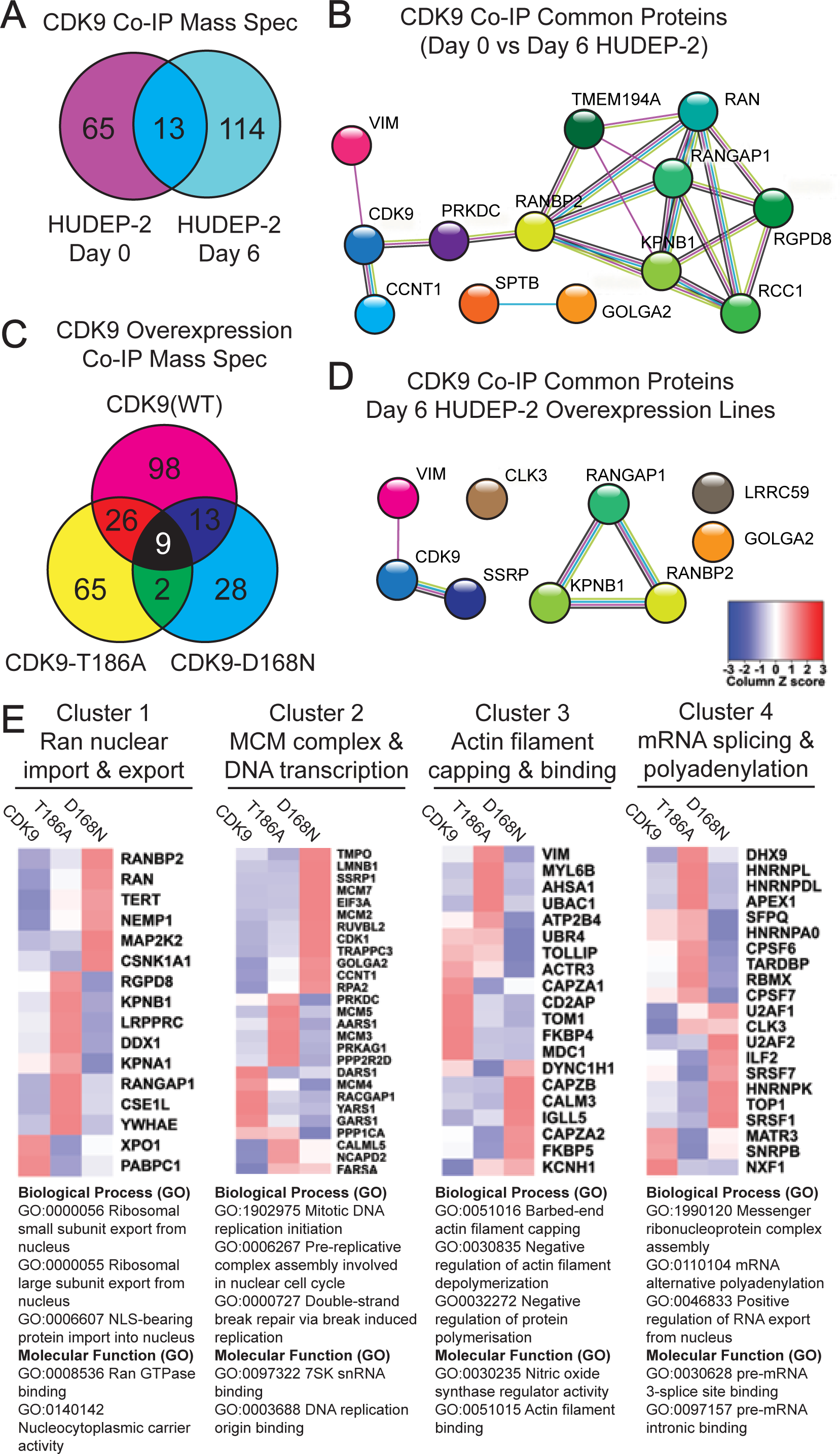
Summary of CDK9 co-immunoprecipitation mass spectrometry results in HUDEP-2 cells. **(A)** Venn diagram of identified interactors of CDK9 in undifferentiated (Day 0) and differentiated (Day 6) HUDEP-2 cells. Proteins were considered an interactor when present in all 3 replicates and ≥3 fold enriched compared to the IgG controls. See the supplementary file for complete list of identified proteins. **(B)** STRING analysis of the 13 common proteins identified in both undifferentiated (Day 0) and differentiated (Day 6) HUDEP-2 cells. STRING analysis was performed using medium confidence (0.4) for interaction score. Aqua lines represent known interactions from curated databases, purple lines represent experimentally determined known interactors. Green, red, and blue lines represent predicted interactors, yellow lines represent text mining identification and black lines represent known co-expression. **(C)** Venn diagram of identified interactors of CDK9 overexpression, CDK9-T186A and CDK9-D168N lines in differentiated (Day 6) HUDEP-2 cells. Proteins were considered an interactor when present in all 3 replicates and ≥3 fold enriched compared to the IgG controls. See the supplementary file for a complete list of identified proteins. **(D)** STRING analysis of the 9 common proteins identified in all 3 HUDEP-2 CDK9 overexpression cell lines. STRING analysis was performed using medium confidence (0.4) for interaction score. Aqua lines represent known interactions from curated databases, purple lines represent experimentally determined known interactors. Green, red, and blue lines represent predicted interactors, yellow lines represent text mining identification and black lines represent known co-expression. **(E)** Heatmaps of clusters 1-4 described in Supplementary Figure 7. Proteins identified in each cluster were compared to the average abundance found in CDK9, CDK9-T186A and CDK9-D168N overexpression HUDEP-2 cell lines. Top biological processes (GO) and molecular functions (GO) identified in STRING are listed below for each cluster (strength >2, false discovery <0.001).

We next explored whether CDK9 activity impacted the interactome by comparing exogenous CDK9, a constitutively active kinase mutant CDK9-T186A, and a dominant negative kinase mutant CDK9-D168N (Dow et al., 2010). We isolated a polyclonal population of transduced cells by FACS **(Supp. Figure 3)** and subsequently performed CDK9 co-immunoprecipitation mass spectrometry in day 6 differentiated cells. With the cut-off described above, we identified a total of 243 proteins, with 98 uniquely identified in the CDK9 expression line, 65 unique to the CDK9-T186A line and 28 unique to the CDK9-D168N line, with 9 proteins identified in all 3 sets **(Figure 4C)**. A STRING analysis of these 9 intersecting proteins identified again 3 members of the Ran GTPase nuclear import complex, including importin-β, confirming our previous results **(Figure 4D)**. We then performed STRING interactome analysis on all 243 identified proteins and clustered the results using k-means clustering to understand the variety of processes and pathways that CDK9 may regulate in differentiating human erythroblasts **(Supp. Figure 4)**. 11 clusters included processes related to Ran-mediated nuclear export and import (Cluster 1), minichromosomal maintenance complex (MCM) and DNA transcription (Cluster 2), actin filament capping and binding (Cluster 3), and as mRNA splicing and polyadenylation (Cluster 4). These four clusters were selected for further analysis based on overall higher abundance in mass spectrometry protein identification **(Figure 4E)**. The abundance of proteins in each cluster were compared across the CDK9, CDK9-T186A and CDK9-D168N datasets **(Figure 4E)**. Within the nuclear trafficking cluster (Cluster 1), importin-β (KPNB1), which drives nuclear import, was enriched in the constitutively active kinase mutant (T186A), but exportin-1, which drives nuclear export, was depleted **(Figure 4E, Panel 1)**. This is compatible with the notion that CDK9 phosphorylation may regulate its nucleocytoplasmic shuttling as has been previously reported (Napolitano et al., 2002).

CDK9 and cyclin T1 were more abundant in the dominant negative (D168N) CDK9 mutant lines, indicative of a relationship between CDK9 phosphorylation state and turnover of CDK9 and cyclin T1 **(Figure 4E, Panel 2)**. Additionally, CDK9, cyclin T1 and other interactors within the minichromosomal maintenance (MCM) complex & DNA transcription cluster (Cluster 2) were enriched in the dominant negative kinase mutant (D168N), suggesting CDK9 kinase activity may impact the expression of various components involved in DNA transcription, including cyclin T1 and CDK9 itself **(Figure 4E, Panel 2)**. Interestingly, CDK9 has been shown to regulate DNA damage responses through MCM complex members through interaction with cyclin K (Yu & Cortez, 2011), which was not detected in our assay. CDK9 was found to interact with several components involved in actin filament capping & binding, which may carry implications for how CDK9 affects actin-related processes during enucleation **(Figure 4E, Panel 3)**. Finally, examination of the mRNA splicing & polyadenylation complex reveals shifts in how CDK9 kinase functionality may impact on its previously reported role in mRNA splicing **(Figure 4E, Panel 4)** (Hu et al., 2021). Here, our identification of a physical association between CDK9 and the Ran-Importin-β nuclear import complex ties together our original finding of an RNA Pol II-independent role for CDK9, and recent findings that NEMP1 regulates nuclear envelope openings during erythroid enucleation.

### Importin-β is required for erythroid enucleation upstream from Calcium signalling and F-actin polymerisation

We next tested the function of importin-β in the context of enucleation using the reversible inhibitor of importin-β-mediated nuclear import, importazole (Soderholm et al., 2011). Inhibition of importin-β using importazole arrested enucleation in a dose-dependent manner in mouse orthochromatic erythroblasts, demonstrating that importin-β function is critical for enucleation **(Figure 5A)**. NEMP1, which has been implicated in maintain nuclear envelope openings which are essential for enucleation (Hodzic et al., 2022), was identified in complex with CDK9 and importin-β, and so we aimed to assess the localisation of importin-β during both normal and arrested enucleation as a comparison. Morphological analysis of orthochromatic erythroblasts treated with importazole indicated an increased number of cells arrested with a polarised nucleus **(Figure 5A)**, phenocopying both CDK9 and F-actin inhibitors and suggesting that importin-β is acting prior to nuclear extrusion. Furthermore, inhibition of importin-β supports the observed phenotype in NEMP1 knockout mice (Hodzic et al., 2022). Additionally, we treated day 12 differentiated HUDEP-2 cells with importazole for 24 hours and counted enucleated cells versus non-enucleated and found a significant and dose-dependent reduction in enucleation, indicating conserved function in human erythroid enucleation **(Figure 5B)**. To test whether importin-β regulated the nucleocytoplasmic import of CDK9, we imaged fixed, undifferentiated HUDEP-2 cells following treatment with NVP-2, importazole or DMSO to assess the nuclear to cytoplasmic ratio of active CDK9, cyclin T1 and importazole. Blocking CDK9 activity did not alter the nucleocytoplasmic ratio of CDK9, cyclin T1 or importin-β **(Supp. Figure 5)**. Interestingly, importazole significantly altered the localisation of cyclin T1, but not CDK9 **(Supp. Figure 5)**. This result, taken together with observed co-immunoprecipitation of CDK9 and importin-β **(Figure 4)**, suggests that importin-β may interact with CDK9 for reasons other than nucleocytoplasmic transport in the context of enucleation.

**Figure 5.**
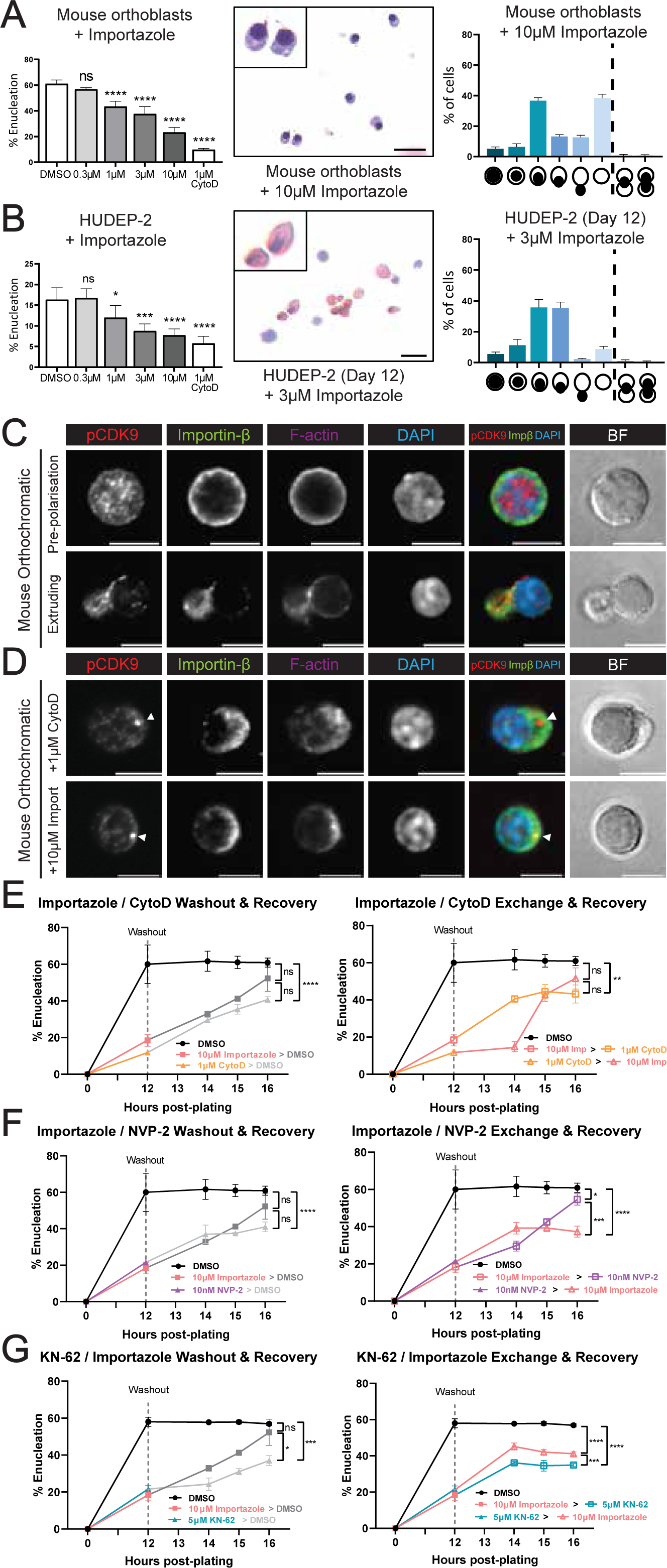
Importin-β activity is required for erythroid enucleation processes downstream of CDK9. **(A)** Quantification of enucleation of mouse orthochromatic erythroblasts following treatment with importazole for 12 hours. DMSO (vehicle control) and cytochalasin D (positive control) are included, in addition to cytospin rapid diff staining and phenotype analysis. n = 4 replicates across 3 independent experiments. (ns = not significant, \*\**p* < 0.01, \*\**p* < 0.01, \*\*\*\**p* < 0.0001; one-way ANOVA with Dunnett’s multiple comparisons test). **(B)** Quantification of enucleation of day 12 differentiated HUDEP-2 cells following treatment with importazole for 12 hours. DMSO (vehicle control) and cytochalasin D (positive control) are included, in addition to cytospin rapid diff staining and phenotype analysis. n = 4 replicates across 3 independent experiments. (ns = not significant, \*\**p* < 0.01, \*\**p* < 0.01, \*\*\*\**p* < 0.0001; one-way ANOVA with Dunnett’s multiple comparisons test). **(C)** Immunofluorescence confocal microscopy of mouse orthochromatic erythroblasts at the pre-polarisation and extrusion phases of erythroid enucleation stained for phospho-CDK9(Thr186), importin-β, phalloidin for F-actin and DAPI for nuclei. Brightfield (BF) and F-actin are excluded from the merge. Scale bars = 5 µm. **(D)** Immunofluorescence confocal microscopy of mouse orthochromatic erythroblasts following 12 hours treatment with 1 μM cytochalasin D or 10 μM importazole, stained for phospho-CDK9(Thr186), importin-β, phalloidin for F-actin and DAPI for nuclei. Brightfield (BF) and F-actin are excluded from the merge. Arrows indicate accumulation of CDK9. All scale bars = 5μm. **(E)** Enucleation rates of mouse orthochromatic erythroblasts following inhibitor washout into fresh media containing DMSO (first plot) or inhibitors (cytochalasin D or importazole; second plot) at 12 hours post-plating. Enucleation rates were measured at 12-, 14-, 15- and 16-hours post-plating. DMSO (black; negative control) is included. Colours change to match either DMSO (grey) or inhibitors (coloured) after washouts. Enucleation rates at hour 16 were statistically compared. (ns = not significant, *\*\*p* < 0.01, \*\*\*\**p* < 0.0001; two-way ANOVA with Tukey’s multiple comparisons test). **(F)** Enucleation rates of mouse orthochromatic erythroblasts following inhibitor washout into fresh media containing DMSO (first plot) or inhibitors (NVP-2 or importazole; second plot) at 12 hours post-plating. Enucleation rates were measured at 12-, 14-, 15- and 16-hours post-plating. DMSO (black; negative control) is included. Colours change to match either DMSO (grey) or inhibitors (coloured) after washouts. Enucleation rates at hour 16 were statistically compared (ns = not significant, *\*\*p* < 0.01, \*\*\*\**p* < 0.0001; two-way ANOVA with Tukey’s multiple comparisons test. **(G)** Enucleation rates of mouse orthochromatic erythroblasts following inhibitor washout into fresh media containing DMSO (first plot) or inhibitors (KN-62 or importazole) at 12 hours post-plating. Enucleation rates were measured at 12-, 14-, 15- and 16-hours post-plating. DMSO (black; negative control) is included. Colours change to match either DMSO (grey) or inhibitors (coloured) after washouts. Enucleation rates at hour 16 were statistically compared (ns = not significant, *p < 0.1, ***p < 0.001; two-way ANOVA with Tukey’s multiple comparisons test).

As importazole appeared to block enucleation at a similar stage to CDK9 and F-actin inhibitors, we assessed which of these proteins might regulate each other during enucleation. We first examined importin-β localisation using fluorescence confocal microscopy. Importin-β appears to localise to the nuclear envelope in pre-polarised erythroblasts with a similar phenotype reported for NEMP1 (Hodzic ref) **(Figure 5C)** and is observed in the cytoplasm and near folded regions at the nuclear envelope, before colocalising with p-CDK9 in the future reticulocyte during nuclear extrusion **(Figure 5C)**. Cells arrested by treatment of either cytochalasin D (CytoD) or importazole resulted in an accumulation of p-CDK9 at the site of enucleation, with importin-β localising around the membrane opposite to the polarised nucleus **(Figure 5D)**. Because p-CDK9 accumulates to the extrusion site in the same way under both actin and importin-β inhibitors, we postulate that p-CDK9, similarly to actin, may be required for enucleation upstream of importin-β before nuclear extrusion.

To test this functionally, we performed the previously described inhibitor washout, exchange, and recovery enucleation assays to determine an order of events using importazole, cytochalasin D (CytoD) and NVP-2. Enucleation rates were assessed by flow cytometry following 12 hours incubation with importazole, cytochalasin D (CytoD) or NVP-2 and compared to the vehicle (DMSO) control directly after washout, and at hour 14, 15 and 16 after plating cells **(Figure 5E, F)**. Enucleation rates in the vehicle (DMSO) treated control group remain steady, and enucleation rates following importazole washout were able to recover, confirming the reversibility of the blockage by importazole **(Figure 5E, Panel 1)**. Inhibition of importin-β (importazole) before F-actin (CytoD) led to an initial recovery of enucleation rates but an overall reduction by hour 16 **(Figure 5E, Panel 2)**. Conversely, blocking F-actin (CytoD) before importin-β (importazole) resulted in a gradual recovery of enucleation rates **(Figure 5E, Panel 2)**. This indicates that importin-β is required at an earlier stage of enucleation than F-actin polymerisation. Blocking importin-β (importazole) before CDK9(NVP-2) resulted in a consistent recovery of enucleation rates, but the reverse sequence did not **(Figure 5F)**. We repeated this by inhibiting CaMKII (KN-62) and importin-β (importazole); however, we did observe altered recovery rates for KN-62 in this experiment, where recovery was significantly reduced compared to the control at hour 16 **(Figure 5G, Panel 1)**. Nonetheless, we could observe a clear shift in recovery when washing out importazole into KN-62, which resulted in a reduction of enucleation, as opposed to washout of KN-62 into importazole which allowed for a partial recovery in enucleation **(Figure 5G, Panel 2)**. Taken together, this supports our previous findings that CaMKII acts upstream of F-actin and indicates CaM signalling is necessary downstream of CDK9 but may occur dynamically or simultaneously to importin-β activity. We conclude that importin-β is essential for enucleation, acting downstream of CDK9 and upstream of F-actin, and that both importin-β and CDK9 are required near the membrane opposite the polarised nucleus prior to nuclear extrusion. A proposed order of the molecular events regulating enucleation is summarised in **Figure 6**, alongside non-mutually exclusive potential models for the role of CDK9 and importin-β in erythroid enucleation.

**Figure 6.**
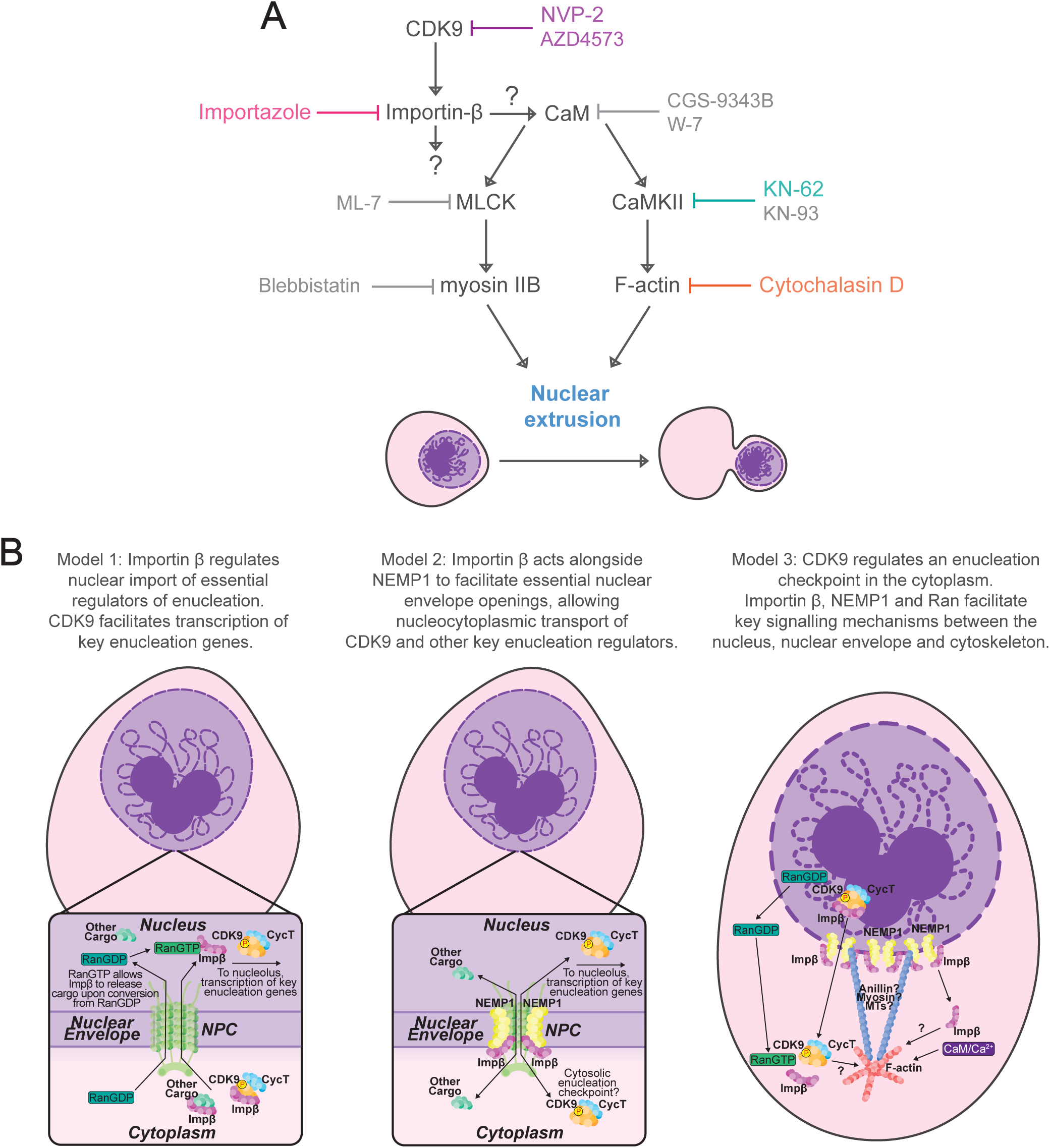
Proposed non-mutually exclusive models for the role of CDK9 and importin-β in regulating erythroid enucleation prior to CaM/Ca^2+^ signalling. **(A)** Order of action model of CDK9 activity to achieve nuclear extrusion by downstream activation of the calmodulin (CaM) pathway through direct, or indirect activity of importin-β. CaM activation results in CaMKII and MLCK activation which in turn results in F-actin polymerisation and myosin IIB contraction respectively to achieve nuclear extrusion. Inhibitors shown in colours correspond to drugs used throughout this study. Inhibitors shown in grey have been previously described in this context (Wölwer et al., 2016). **(B)** Non-mutually exclusive models depicting potential roles for CDK9 and importin-β in enucleation. NPC = Nuclear pore complex. Model 1: Importin-β regulates nuclear import of essential regulators of enucleation. CDK9 facilitates transcription of key enucleation genes. Model 2: Importin-β acts alongside NEMP1 to facilitate essential nuclear envelope openings, allowing nucleocytoplasmic transport of CDK9 and other key enucleation regulators. Model 3: CDK9 regulates an enucleation checkpoint in the cytoplasm. Importin-β, NEMP1 and Ran facilitate key signalling mechanisms between the nucleus, nuclear envelope and cytoskeleton.

## Discussion

### CDK9 and importin-β facilitate cellular reorganisation prior to CaM/Ca^2+^ signalling and F-activity to achieve enucleation

Our previous work had identified CDK9 as a regulator of enucleation (Wölwer et al., 2015), and this study aimed to delineate the mechanisms involved. In doing so, we have identified importin-β as a novel regulator of enucleation and direct interactor of CDK9. How may importin-β and CDK9 coordinate erythroid enucleation? Here, we propose three non-mutually exclusive models **(Figure 6B)**. The typical biological function of CDK9 is well-characterised, where it acts as the kinase component and small subunit of P-TEFb to regulate RNA polymerase II (RNA Pol II) transcriptional elongation by phosphorylating the carboxyl terminal domain (CTD) of RNA Pol II and negative regulators DRB sensitivity-inducing factor (DSIF) and negative elongation factor (NELF) (Anshabo et al., 2021; Gilchrist et al., 2008; Paparidis et al., 2017; Wada et al., 1998). Our previous study had shown that blocking the activity of RNA Pol II in late erythropoiesis does not arrest enucleation (Wölwer et al., 2015), suggesting a new role for CDK9 in enucleation independently of RNA Pol II. Neither DSIF or NELF have been shown to interact with or modulate the activity of anything other than RNA Pol II and we did not identify them in our proteomics search, therefore we did not investigate their potential role in enucleation but at this stage cannot rule out a possible involvement (Aoi et al., 2020; Decker, 2021; Deng et al., 2022). Considering the localisation of CDK9 during enucleation and the phenotype of CDK9 inhibitor-induced arrest, we conclude that CDK9 must be acting within the cytoplasm prior to nuclear extrusion, however it remains plausible that CDK9-mediated transcription via RNA Pol II is essential for enucleation, and importin-β may play a role in the nucleocytoplasmic location of CDK9 **(Figure 6B: Model 1)**.

NEMP1 plays an essential role in enucleation by modulating nuclear envelope openings that are essential for enucleation, and NEMP1 knockout mice show anaemia due to apoptosis of polychromatic erythroblasts (Hodzic et al., 2022). Considering both NEMP1 and importin-β are key components of the Ran nuclear import pathway (Shibano et al., 2015), it is likely that the function of importin-β in enucleation is related to that of NEMP1. Interestingly, NEMP1 has been shown to support the mechanical stiffness of the nuclear envelope in some settings (Tsatskis et al., 2020), potentially implicating NEMP1 (and by association importin-β) in the mechanical processes involved in extrusion of the nucleus during enucleation. Considering CDK9 and importin-β were found to physically interact with NEMP1 in the context of erythropoiesis, we postulate that importin-β, NEMP1 and possibly other members of the Ran nucleocytoplasmic import complex may regulate changes to the nuclear envelope to assist in the shuttling of key enucleation regulators, including CDK9, between the cytoplasm and nucleus prior to nuclear extrusion **(Figure 6B, Model 2)**.

Finally, our previous work has identified a role for calcium signalling in enucleation via the calmodulin (CaM) pathway, and the role of F-actin is well documented (Newton et al., 2024; Wölwer et al., 2016). More recently, Zhang et al (2015) describe and support the notion of finely tuned CaM/Ca^2+^ signalling during erythropoiesis and enucleation (Zhang et al., 2025). Here, we examined the order of which CDK9 and importin-β may act to regulate enucleation in respect to calcium signalling and F-actin activity and found both to function upstream. The calmodulin (CaM) pathway is implicated in both myosin and F-actin activation, through the activity of myosin light chain kinase (MLCK) and Ca^2+^/CaM-dependent kinase II (CaMKII) respectively (Mizuno et al., 2008; Okamoto et al., 2007). Calmodulin signalling has been shown to regulate CDK9 T-loop phosphorylation, which is required for its activation, which may suggest a coordinated role for calmodulin signalling in separate steps in enucleation involving CDK9 and the cytoskeleton (Ramakrishnan & Rice, 2012). Additionally, calcium signalling, and intracellular calcium levels have been shown to block nuclear import by altering the activity and localisation of RanGTP and importin-β (Kaur et al., 2014; Sweitzer & Hanover, 1996). Strikingly, blocking enucleation with an actin inhibitor or importin-β inhibitor both caused a reproducible and consistent accumulation of phosphorylated CDK9 near the junction of enucleation. This may be due to the aggregation of activated CDK9 at a critical site of activity, which we speculate may be the proposed ‘enucleosome’ (Nowak et al., 2017), therefore priming the cells for nuclear extrusion following release of F-actin or importin-β inhibition. We propose that CDK9 and importin-β may function as part of a checkpoint in the final stages of enucleation, in conjunction with an influx of calcium and subsequent calmodulin (CaM) signalling to achieve nuclear extrusion **(Figure 6B, Model 3)**.

### Characterisation of the CDK9 interactome in terminally differentiating erythroblasts

This study provides new insights into the CDK9 interactome during erythroblast development. Importantly, we identified several components of the Ran nuclear import pathway, including importin-β and RanGTP itself, which are associated directly with CDK9 in both undifferentiated and differentiated HUDEP-2 cells. How might RanGTP act to regulate enucleation? RanGTP acts as a master controller for nucleocytoplasmic transport through the nuclear pore complex (NPC) by interactions with importins and exportins (Cavazza & Vernos, 2016).importin-β. CDK9 is known to shuttle between the nucleus and cytoplasm, and other interactors of CDK9 including cyclin T and HEXIM have been shown to interact with importin α to facilitate transport through the nuclear pore complex (Czudnochowski et al., 2010; Napolitano et al., 2002). CDK9 has previously been shown to interact with nuclear pore components RANGAP1, RANBP2 and RGPD3 (Ramakrishnan et al., 2012), which we also identified as part of the RanGTP complex that interacts with CDK9 in erythroid cells. However, the previous study only focused on localisation of CDK9-Cyclin T1 to the nuclear pore and did not examine the role of Ran and importin-β in CDK9-Cyclin T1 nuclear import. Of note, exportin 7 (Xpo7), which can facilitate nuclear export through the RanGTP pathway, is essential for histone export into the cytoplasm during the nuclear condensation phase of enucleation (Hattangadi et al., 2014; Mingot et al., 2004). importin-βimportin-βIt will be interesting to examine the relationship between the RanGTP pathway and exportin 7 (Xpo7) in the control of histone export and how CDK9 may be involved in this process. While this study offers novel insights into the potential regulation of CDK9 and cyclin T1 nucleocytoplasmic transport via importin-β interactions,, we have clearly shown that importin-β regulates enucleation by direct interaction with CDK9 and NEMP-1 in erythroblasts and is required upstream of CaM/Ca^2+^ signalling and F-actin activity.

What else might the CDK9 and importin-β interactome tell us about enucleation? Besides nuclear import, importin-β has been shown to modulate the permeability and structure of the nuclear pore complex (NPC) through direct interaction with nuclear pore complex protein Nup153 (Lowe et al., 2015), highlighting a potential role in modifying nuclear structure to facilitate enucleation. Additionally, importin-β is implicated in wider transport roles, where it can regulate kinesin-4 motor activity in plant cells (Ganguly et al., 2018), and can directly bind to certain cargo, such as the parathyroid hormone (PTH)-related protein (PTHrP), to facilitate microtubule-dependent transport (Lam et al., 2002). Importantly, importin-β is implicated in the regulation of anillin during cytokinesis (Beaudet et al., 2020); anillin acts to scaffold components of the actomyosin ring, including F-actin, myosin and RhoA (Piekny & Glotzer, 2008), all of which are essential for nuclear extrusion during enucleation. Examining whether anillin is required for enucleation and how this may relate to importin-β function will be key to investigating this link. Finally, we observed nuclear envelope protein 1 (NEMP1/TMEM194A) across all sample sets, which forms part of the Ran nuclear import pathway, by recruiting Ran to the nuclear envelope (Shibano et al., 2015).

In addition to the Ran import pathway complex, we also identified several other new interactors of CDK9. Golgin A2 (GOLGA2), which was observed across all sample sets and was enriched in CDK9 pulldowns, acts to maintain Golgi structure through stacking of Golgi cisternae and has been investigated as a therapeutic target in cancer due to its relationship with autophagy (Chang et al., 2012; Nakamura et al., 1995). As autophagy is essential for erythroid terminal differentiation, it will be important to examine the relationship between CDK9 and golgin A2-mediated autophagy mechanisms during enucleation. Vimentin (VIM), which is an intermediate filament protein implicated in the function of many cells types (Danielsson et al., 2018) was also identified across all sample sets, and has been previously identified in a similar CDK9 immunoprecipitation mass spectrometry interactome set (Yang et al., 2015). Yang et al. (2015) identified a CDK9 interactome in human A549 pulmonary epithelial cells, and some of these proteins are also identified in our data (Yang et al., 2015). However, most proteins in our identified interactome are new, likely reflecting the specialised role CDK9 may play in erythroid development.

Of note, we identified a large complex involved in minichromosome maintenance (MCM) and DNA damage response. Although CDK9 has been shown to play a role in maintaining genome integrity in partnership with cyclin K (Yu & Cortez, 2011), we did not detect cyclin K in our assay, and it is therefore possible that this mechanism extends to erythroblasts through a different cyclin. This should be investigated further. Furthermore, a role for CDK9 in mRNA splicing has been described (Hu et al., 2021), and with the identification of mRNA splicing mechanisms, polyadenylation factors and several members of the messenger ribonucleoprotein complex assembly, also known as mRNP granules (Buchan, 2014) in human erythroblasts, new insights could be gained on the role of CDK9 in mRNA processing in erythropoiesis. Importantly, we also identified a large complex involved in actin filament capping and binding, including CAPZA1/2, CAPZB, amongst others, which may point to how CDK9 might interact with actin during enucleation processes. Additionally, we identified tropomodulin-1, which has been observed to co-localise with F-actin and non-musical myosin IIB during enucleation (Nowak et al., 2017), and Cdc42, which regulates essential polarisation of the nucleus prior to nuclear extrusion (Ubukawa et al., 2020). The relationship between CDK9 and the regulation of actinomyosin polymerisation should be investigated further, using genetic models for proximity labelling and functional assays to determine how CDK9 activity may be involved in modulating actinomyosin activity in the context of erythroid terminal differentiation. Although we don’t fully understand how CDK9, importin-β and downstream effectors of enucleation directly interact to coordinate enucleation, our studies provide insight into a series of events that must occur before F-actin mediated nuclear extrusion and presents possible new interactors of CDK9 and importin-β in the context of erythropoiesis and erythroid enucleation.

## Acknowledgements

This work was completed on Wurundjeri land of the Kulin nation, and we pay our respects to the Elders past and present. We acknowledge Dr. Margaret Veale and the La Trobe Bioimaging Platform for flow cytometry and optical microscopy. We acknowledge Dr. Pierre Faou, Dr. Rohan Lowe, Dr. Keshava Datta and the La Trobe Proteomics and Metabolomics Platform for mass spectrometry and proteomics. We acknowledge the La Trobe Animal Research & Training Facility (LARTF) for supporting animal work. We thank Dr. Ryo Kurita from the Riken BioResource Research Center (Tsukuba, Japan) for providing HUDEP-2 cell lines, and Prof. Jan Frayne from the University of Bristol (Bristol, United Kingdom) for providing BEL-A cell lines. LMN was supported by an Australian Government Research Training Program (RTP) scholarship. POH was supported by NHMRC Fellowship APP1079133. This work was supported by NHMRC Project grant APP1103858 (POH) and ARC Discovery project grant DP190103634 (POH and EDH).

## Authorship

### Contributions

POH conceived of the project. LMN designed and performed experiments. LMN, KYBL, DYA, CBW, EDH and POH contributed to data analysis and optimisation of experimental protocols. SMR provided resources. CJ contributed to image acquisition and analysis. LMN, SMR and POH drafted the manuscript. All authors contributed to editing the manuscript. EDH and POH supervised the study.

### Correspondence

Patrick Humbert, Department of Biochemistry and Chemistry, La Trobe Institute for Molecular Science, La Trobe University, Kingsbury Drive, Bundoora, Victoria, Australia, 3073.

## Supplementary Figure Legends

**Supplementary Figure 1.**
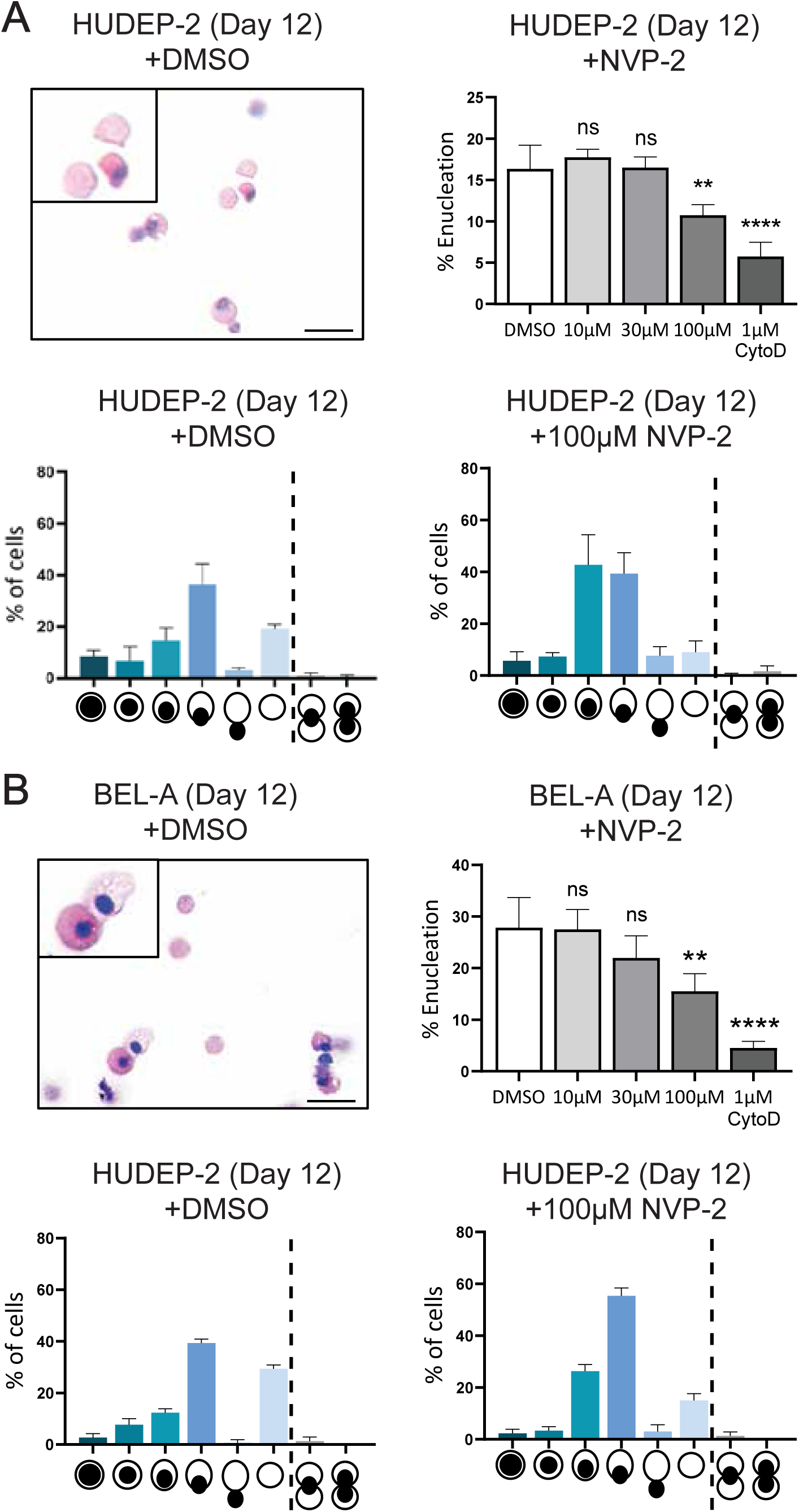
NVP-2 blocks enucleation of HUDEP-2 and BEL-A cells. **(A)** Quantification of enucleation of differentiated HUDEP-2 cells (Day 12) following 24-hour treatment with NVP-2. DMSO (vehicle control) and cytochalasin D (positive control) are included, in addition to cytospin rapid diff staining of DMSO control cells and phenotype analysis of DMSO control and NVP-2 treated cells. n = 205 cells counted (DMSO) and 186 cells counted (NVP-2). (ns = not significant, \*\**p* < 0.01, \*\*\*\**p* < 0.0001; one-way ANOVA with Dunnett’s multiple comparisons test). Scale bar = 10 µm. **(B)** Quantification of enucleation of differentiated BEL-A cells (Day 12) following 24-hour treatment with NVP-2. DMSO (vehicle control) and cytochalasin D (positive control) are included, in addition to cytospin rapid diff staining of DMSO control cells and phenotype analysis of DMSO control and NVP-2 treated cells. n = 235 cells counted (DMSO) and 168 cells counted (NVP-2). (ns = not significant, \*\**p* < 0.01, \*\*\*\**p* < 0.0001; one-way ANOVA with Dunnett’s multiple comparisons test). Scale bar = 10 µm.

**Supplementary Figure 2.**
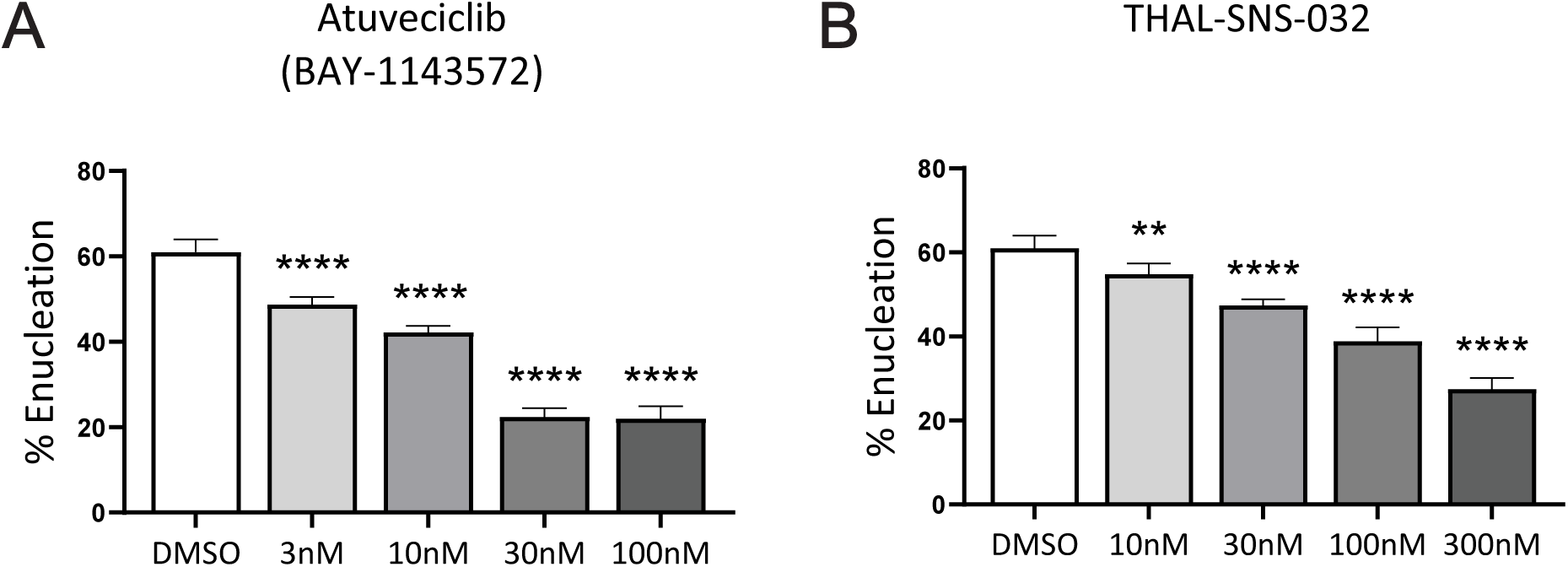
CDK9 inhibitors Atuveciclib and THAL-SNS-032 cause a dose dependent arrest of mouse erythroblast enucleation. **(A)** Quantification of enucleation of mouse orthochromatic erythroblasts following treatment with Atuveciclib (BAY1143572) for 12 hours. DMSO (vehicle control) is included. n = 4 replicates across 2 independent experiments. (\*\*\*\**p* < 0.0001; one-way ANOVA with Dunnett’s multiple comparisons test). **(B)** Quantification of enucleation of mouse orthochromatic erythroblasts following treatment with THAL-SNS-032 for 12 hours. DMSO (vehicle control) is included. n = 4 replicates across 2 independent experiments. (\*\**p* < 0.01, \*\*\*\**p* < 0.0001; one-way ANOVA with Dunnett’s multiple comparisons test).

**Supplementary Figure 3.**
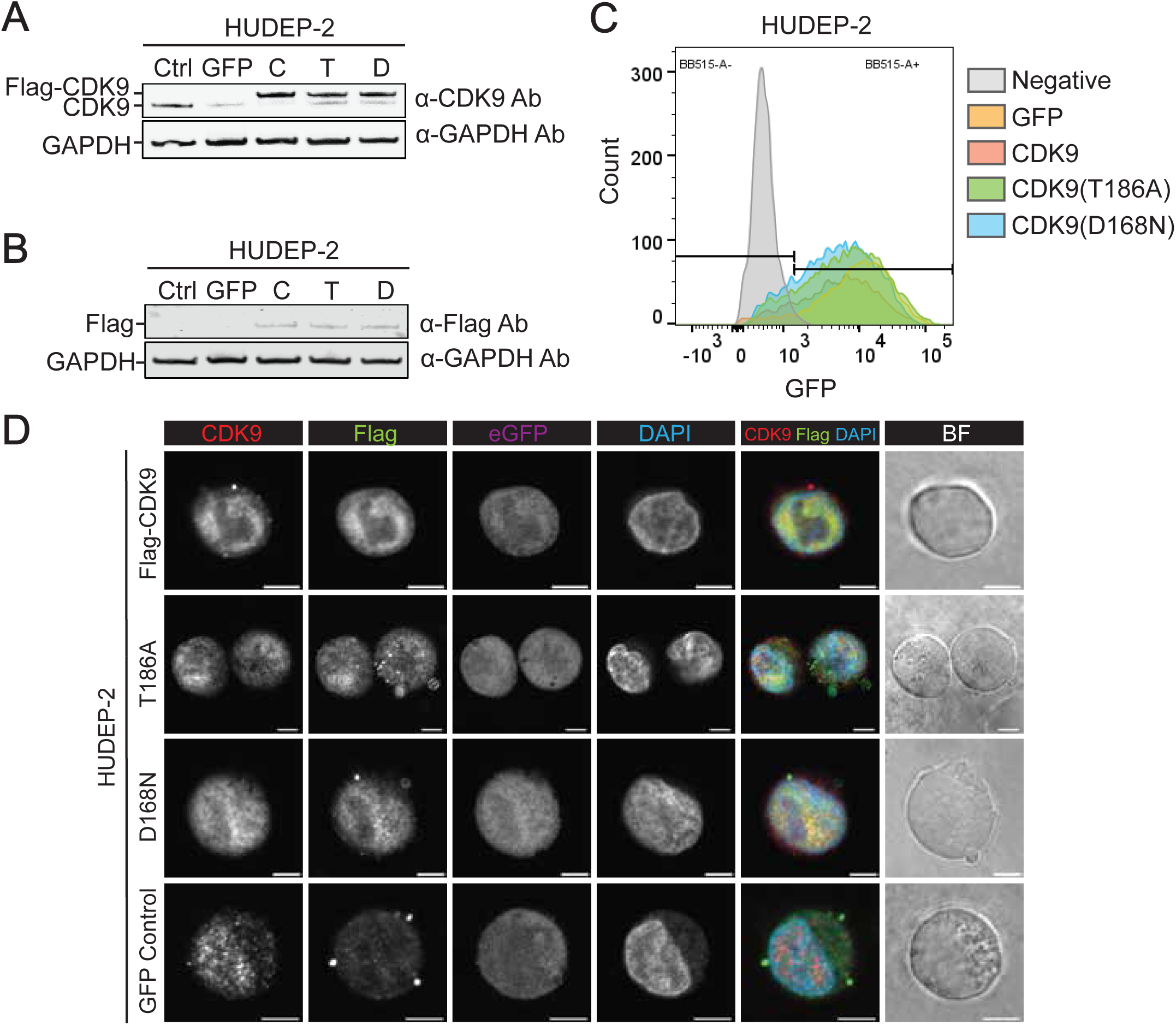
Characterisation of HUDEP-2 CDK9 overexpression cell lines. **(A)** Western blot of HUDEP-2 cell lines, including the GFP vector control, CDK9, CDK9-T186A and CDK9-D168N overexpression lines. Bands can be detected for Flag-CDK9 in the three CDK9 overexpression lines. GAPDH is shown as a loading control. **(B)** Western blot of HUDEP-2 cell lines, including the GFP vector control, CDK9, CDK9-T186A and CDK9-D168N overexpression lines. Bands can be detected for Flag in the three CDK9 overexpression lines. GAPDH is shown as a loading control. **(C)** Flow cytometry histogram showing GFP expression in HUDEP-2 cell lines, including the GFP vector control, CDK9, CDK9-T186A and CDK9-D168N overexpression lines compared to negative control HUDEP-2 cells. Propidium iodide was used to gate for only viable cells. **(D)** Immunofluorescence confocal microscopy of undifferentiated HUDEP-2 cells transduced to overexpress CDK9, CDK9-T186A, CDK9-D168N or GFP only and stained for CDK9, Flag, GFP and DAPI for nuclei. Transduced cells expressing low levels of eGFP are shown in comparison to the wildtype. Brightfield (BF) and eGFP are excluded from the merge. Yellow colour depicts co-localisation of CDK9 and Flag. All scale bars = 5 μm.

**Supplementary Figure 4.**
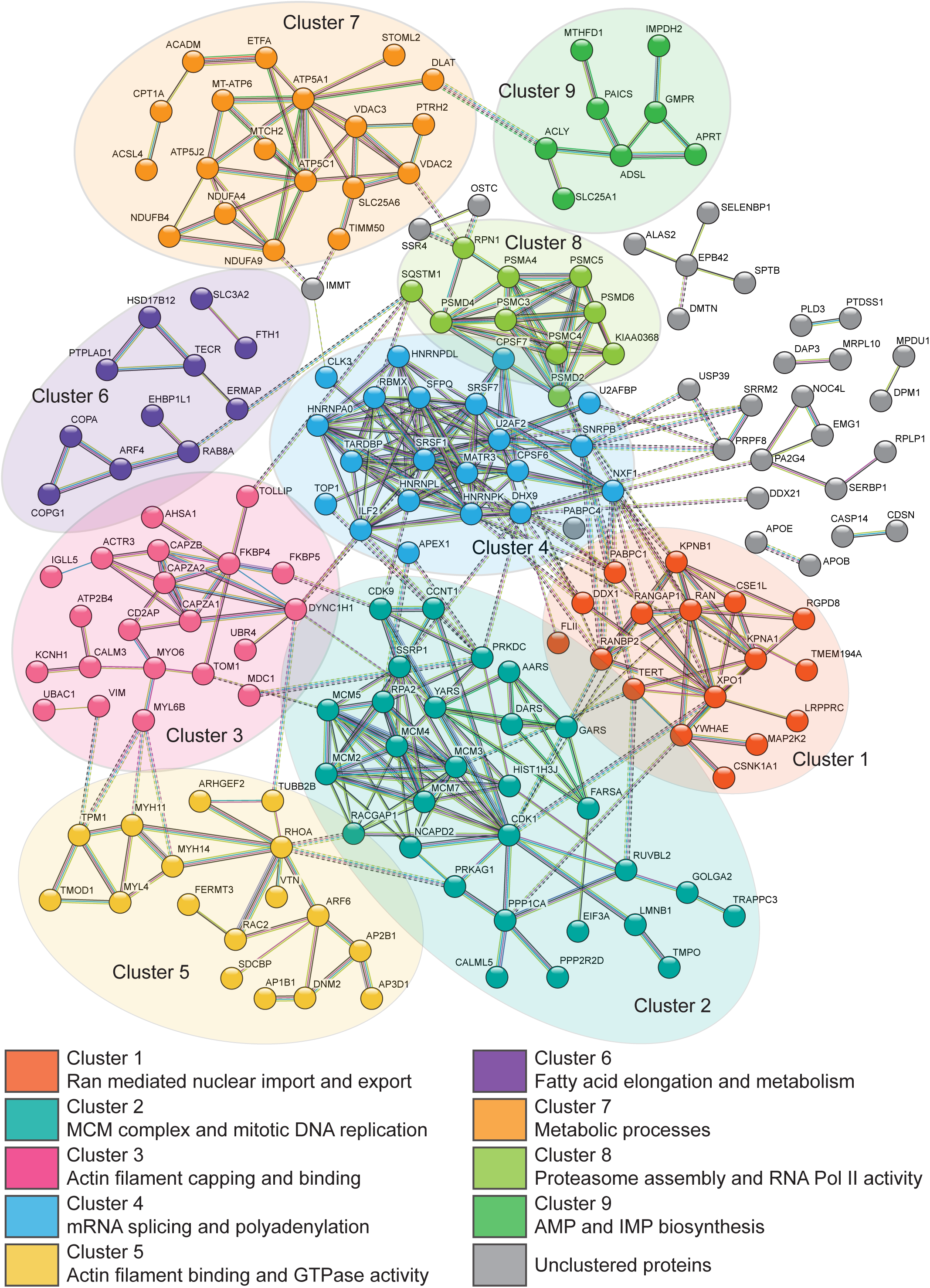
Interactome of identified proteins in HUDEP-2 CDK9 overexpression cell lines. STRING analysis of proteins identified across all HUDEP-2 overexpression lines. Proteins were considered an interactor of CDK9 if identified in all 3 replicates and were ≥3 fold enriched compared to the IgG control. Clusters were identified using STRING analysis of all proteins identified in overexpression lines (243 total proteins), with high confidence (0.7) for interaction score, with unconnected nodes removed. Clustering was performed using kmeans clustering into 11 groups. Aqua lines represent known interactions from curated databases, purple lines represent experimentally determined known interactors. Green, red, and blue lines represent predicted interactors, yellow lines represent text mining identification and black lines represent known co-expression. Dotted lines represent connections between clusters. See the supplementary file for a complete list of identified proteins and corresponding abundances.

**Supplementary Figure 5.**
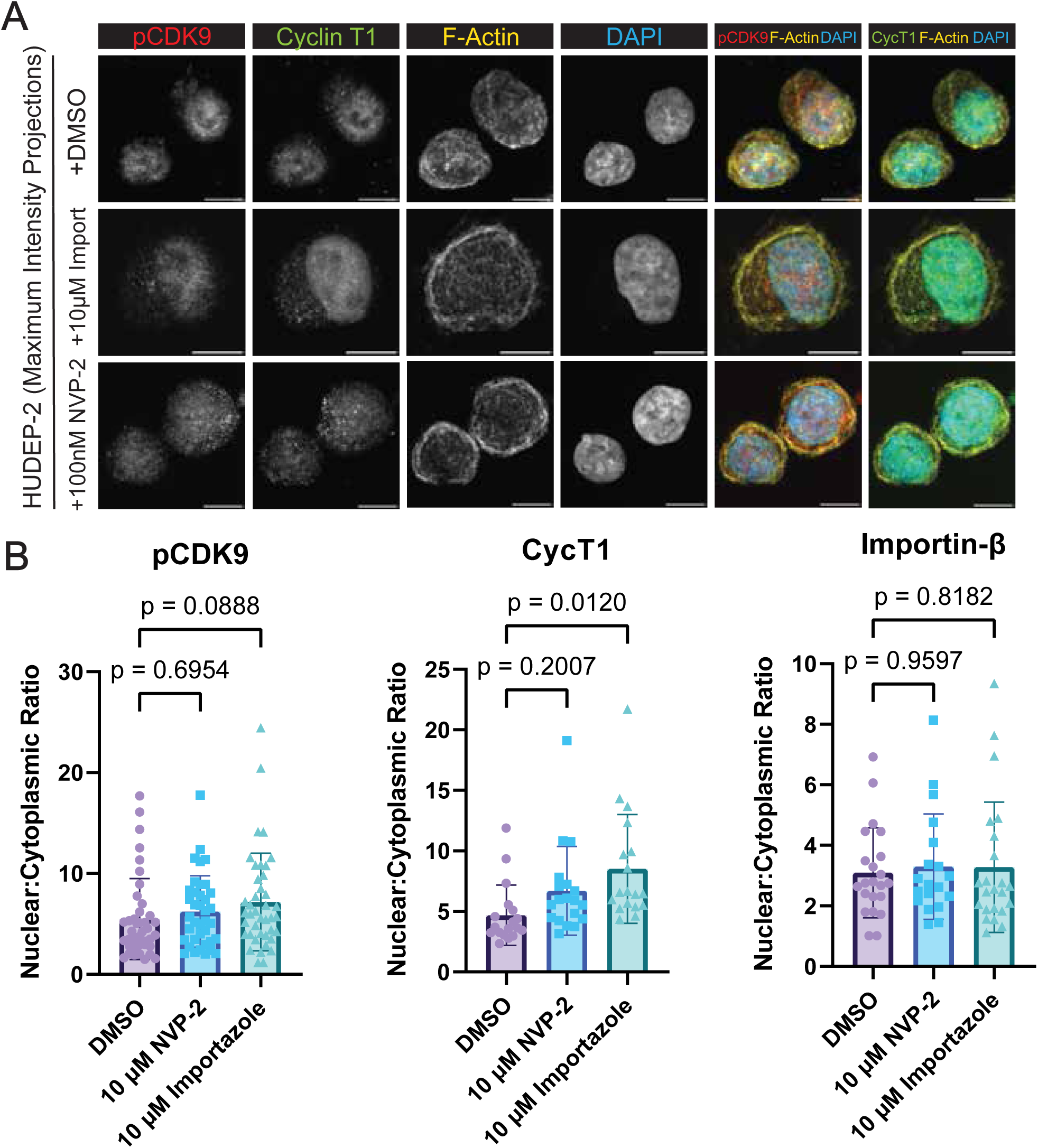
Importazole alters the nucleocytoplasmic ratio of Cyclin T1 but not phosphorylated CDK9. **(A)** Immunofluorescence confocal microscopy of undifferentiated (Day 0) HUDEP-2 cells stained for phospho-CDK9(Thr186), cyclin T1, F-actin and DAPI. Merges showing pCDK9 and cyclin T1 alongside F-actin are shown. All scale bars = 5 μm. **(B)** Analysis of nuclear to cytoplasmic ratio of pCDK9, cyclin T1 and importin-β in fixed undifferentiated (Day 0) HUDEP-2 cells following overnight treatment with 10µM NVP-2, 10µM Importazole or DMSO vehicle control. Each point represents an individual cell. pCDK9 DMSO n = 40, NVP-2 n = 38, Importazole n = 41; CycT1 DMSO n = 18, NVP-2 n = 19, Importazole n = 19; Importin-β DMSO n = 22, NVP-2 n = 19, Importazole n = 23. (p values are shown for each comparison; one-way ANOVA with Dunnett’s multiple comparisons test).

