## Supplemental Methods for "CDK9 interacts with a RanGTP-NEMP1-Importin-β complex to regulate erythroid enucleation"

**Newton et al (2025) – Supplemental Methods**

*CDK9 co-immunoprecipitation*

Co-immunoprecipitation studies were performed using the Pierce Classic Magnetic IP/Co-IP Kit (Thermo Fisher; Waltham, MA) as per the manufacturer’s instructions. Modifications were as follows; cells were lysed in undifferentiated state or on day 6 of differentiation in NETN buffer (100 mM NaCl, 20 mM Tris-Cl (pH 8.0), 0.5 mM EDTA (pH 8.0) and 0.5% (v/v) NP-40) supplemented with Complete Mini protease inhibitor and PhosSTOP protease inhibitor as per the manufacturer’s guidelines (Roche; Basel, Switzerland). Protein extracts were quantified using the DC Protein Assay (BioRad; Hercules, CA). Three separate replicate immune complexes were prepared by combining 10 μg of CDK9 (F-6) or 10 μg mouse IgG isotype control antibody with 1mg of cell lysate and incubated while mixing at 4°C overnight. Immunoprecipitation was performed overnight while mixing at 4°C using 0.3 mg of freshly washed A/G magnetic beads. On-bead trypsin digestion was carried out as previously described (Antonicka et al., 2020). Briefly, beads were washed 3 times and resuspended in 20mM ammonium bicarbonate (pH 8) before addition of Tris (2-carboxyethyl) phosphine to a final concentration of 5 mM and incubated at 45°C for 30 minutes to achieve reduction of cysteine disulphide bonds. Iodoacetamide was added to a final concentration of 20 mM and incubated in the dark for 10 minutes to achieve alkylation. Beads were pelleted and resuspended in 20 mM ammonium bicarbonate (pH 8) with 2.5 µg/mL of trypsin before incubation at 37°C overnight. Beads were pelleted and the supernatant collected, before rinsing the beads in 20mM ammonium bicarbonate (pH 8) and recombining with the supernatant. Samples were dried in a centrifugal evaporator and desalted before liquid chromatography tandem mass spectrometry (LC-MS/MS).

*Liquid chromatography and mass spectrometry analysis*

The tryptic peptides were separated on a Thermo Ultimate 3000 RSLC nano UHPLC system and analysed on a Thermo Q-Exactive HF Orbitrap mass-spectrometer (Thermo Fisher Scientific, Waltham, MA). Peptides were loaded onto a PepMap C18 5 µm 2 cm trapping column (Thermo Fisher Scientific, Waltham, MA) and washed at 5 µL/min for 6 min using Buffer C (0.1% (v/v) trifluoracetic acid, 2% (v/v) ACN) before switching the pre-column in line with the analytical column held at 55°C (nanoEase M/Z Peptide BEH C18 Column, 1.7 µm, 130 Å and 75 µm ID × 25 cm; Waters Corporation, Milford, MA). The separation of peptides was performed at 250 nL/min using a linear gradient of buffer A (0.1% (v/v) formic acid, 2% (v/v) ACN) and buffer B (0.1% (v/v) formic acid, 80% (v/v) ACN), starting at 12% buffer B to 30% over 54 min, then rising to 50% B over 10 min followed by 95% B in 6 min. The column was then cleaned for 4 min at 95% B and then percentage of B was brought down to 2% over 5 minutes. The column was equilibrated with 2% B for 15 minutes. Blanks were run between sample injections. MS data were collected in the Data Dependent Acquisition (DDA) mode over a period of 90 minutes. MS1 scan parameters were: 60,000 resolution, m/z range of 350–1500, AGC target 3e6, maximum ion injection time 30 ms. The top seven ions were fragmented per cycle using a normalized collision energy of 28 and dynamic exclusion was carried out for 25s. The isolation window of the quadrupole for precursor isolation was 1.4 m/z. MS2 scan parameters were: 60,000 resolution, AGC target 1e5, maximum ion injection time 110 ms. Lock mass was set to 445.1200 for internal mass calibration.

*Protein database search and analysis*

The protein database searches were conducted using Sequest HT search engine via the Proteome Discoverer 2.4 software suite (Thermo Fisher Scientific, Waltham, MA). Spectra were matched against the Homo sapiens reference proteome downloaded from Uniprot (The UniProt Consortium, 2023). Precursor tolerance was set to 20 ppm and fragment tolerance to 0.05 Da. Two missed trypsin cleavages were permitted. The included static modification was carbamidomethyl of C, and dynamic modifications were oxidation of M, acetylation of protein N-terminus, and deamidation of N/Q. Validation FDR thresholds were 0.01 for strict and 0.05 for relaxed criteria at PSM, peptide and protein levels. Label-free quantification was performed on precursor ion area. Proteins that were present in all three replicates with a minimum 3-fold enrichment compared to the IgG controls were considered relevant. Results, including generation of Venn diagrams and heatmaps, were analysed using FunRich v3.1.4 (Fonseka et al., 2021). Interaction and gene ontology (GO) analysis was performed using the STRING database (Szklarczyk et al., 2023).
